# Elevated α-synuclein attenuates phagocytosis in *SNCA* triplication human iPSC-derived neuron:microglia co-cultures

**DOI:** 10.1101/2022.11.15.516591

**Authors:** Richard Lieberman, Khaled Elnaggar, Kimberly Jesseman, Sarah DeFrancisco, Kelsey Degouveia, Emma Suneby, Hao Wu, L. Alejandro Rojas, John D. Graef

**Affiliations:** Fulcrum Therapeutics, 26 Landsdowne St, 5^th^ floor, Cambridge, MA 02139

## Abstract

Synucleinopathies such as Parkinson’s disease (PD) are characterized by pathologic production, aggregation, and cell-to-cell transmission of α-synuclein (α-syn) protein that results in impaired cellular function. While neurons of the substantia nigra pars compacta express high levels of α-synuclein and are highly vulnerable to its aberrant expression or conformation, brain-resident macrophages (microglia) are also sensitive to abnormal α-synuclein, with recent reports indicating that elevated levels impair phagocytic ability *in vivo* and *in vitro*. To explore the impact of elevated α-syn on microglial function we employed a co-culture model containing iPSC-derived neurons and microglia-like cells. iPSCs from healthy control donors and a Parkinson’s donor with an allelic triplication of the *SNCA* gene locus were differentiated into neurons and microglia-like cells. In monoculture, neurons and microglia generated from the *SNCA* triplication donor expressed higher levels of *SNCA* transcript and protein. Neurons were found to have significantly greater expression of *SNCA* compared to microglia, regardless of donor genotype. Co-cultures of neurons and microglia revealed that microglia cultured with *SNCA* triplication neurons displayed reduction in phagocytosis of fluorescent *E. coli*, irrespective of microglia donor genotype. *SNCA* mRNA and protein expression could be reduced with treatment with an antisense oligonucleotide (ASO) targeting *SNCA*. ASO treatment partially rescued microglia phagocytosis in *SNCA* triplication co-cultures and in co-cultures containing *SNCA* triplication neurons and healthy control microglia. Our results complement and extend previous findings of impaired microglial function in the presence of elevated α-synuclein in a novel patient-derived co-culture model that utilizes more disease-relevant conditions rather than the relaying on the addition of exogenous α-synuclein.

## Introduction

A hallmark of Parkinson’s disease (PD) is the formation of intracytoplasmic inclusions composed of α-synuclein (α-syn, encoded by the *SNCA* gene) (1, 2). While most PD cases are sporadic in nature, monogenic mutations are implicated in about 3-5% of cases (3). Mutations within the *SNCA* gene locus itself, including point mutations, duplications, and triplications, increase the likelihood of formation of pathogenic α-syn species and associate with PD age of onset and severity (4–7). Human genetic studies show a gene-dosage effect of *SNCA* on disease progression, with *SNCA* gene triplication being associated with an early-onset, severe form of PD (6), while duplication of the gene is associated with later onset and clinical phenotypes that resemble idiopathic PD (7, 8). These findings suggest that there is a direct relationship between *SNCA* gene expression and PD progression, and that reduction of *SNCA* could be a therapeutic strategy to curb disease severity. Although mutations within *SNCA* are rare, explaining disease in ~2.5% of unrelated affected carriers (9), sporadic PD cases, or those caused by mutations outside of the *SNCA* gene, are also characterized by aberrant α-syn protein misfolding and accumulation (10), suggesting that α-syn pathology is a common component irrespective of genetic contribution (11), and that reduction of *SNCA* might be beneficial for curbing disease progression in both idiopathic and genetically defined cases of PD.

α-syn is highly expressed in dopaminergic neurons within the substantia nigra pars compacta, and the degeneration of these cells is a defining characteristic of PD. However, other neural cell types, such as microglia, express α-syn and are affected by aberrant levels of the protein, although the mechanisms underlying their contribution to the disease state are still being elucidated. Microglia are brain-resident macrophages involved in numerous homeostatic functions including phagocytosis; the clearance of dead cells, cellular debris, and aggregated proteins (12). The phagocytic competence of microglia/macrophages has been investigated in the context of Parkinson’s disease using primary murine microglia and cell lines, as well as post-mortem human samples and peripheral blood-derived macrophages (for detailed review, see (13)). Recent reports have suggested that excess α-syn negatively impacts the phagocytic competence of microglia (14–16). However, Janda and colleagues (2018) note that while perturbed phagocytosis is observed in these models, it is still inconclusive whether phagocytosis is pathologically activated or defective in Parkinson’s disease (13). Cellular models that more closely recapitulate the human disease may help clarify the impact of aberrant α-syn on microglial phagocytosis and better define its role in PD pathology.

Bi-directional interactions between microglia and neurons are essential to the normal development of the brain (17–19). In vitro, neurons and microglia in co-culture produce a transcriptional and inflammatory profile that resembles more mature and less disease-associated conditions compared to microglia in monoculture (20, 21). Therefore, a cell model that contains multiple cell types relevant to PD pathology might better recapitulate the disease environment and provide further insights into the disease state, particularly if the cells are of human origin. Human fibroblasts can be re-programmed into pluripotent stem cells (22) that can differentiate into various lineages of the central nervous system (23). Protocols have been published that define the conditions to rapidly generate electrophysiologically active excitatory neurons (24) and phagocytically competent microglia-like cells that resemble, at the transcriptomic level, fetal human microglia (20, 25–27), suggesting that human iPSCs provide a tool to investigate α-syn pathology in disease-relevant cell types. In this investigation, we generated excitatory neurons and microglia-like cells from a donor with an *SNCA* locus triplication and compared them to cells derived from an apparently healthy control donor. Co-cultures of neurons and microglia were assembled and used to explore the effects of elevated α -synuclein on phagocytosis. The findings of this study complement prior works and extend our knowledge of phenotypes uncovered in PD patient-derived *in vitro* models.

## Materials and methods

### iPSC culture

Healthy wild type control (WT) iPSCs derived were obtained from Thermo-Fisher Scientific (WT, GIBCO human episomal iPSC line, cat # A18945), which were generated via reprogramming of cord blood derived CD34+ progenitors. Parkinson’s patient derived *SNCA* triplication iPSCs (*SNCA*) were obtained from the NINDS Human Cell and Data Repository (Cell line ID: ND50040), which were derived via the reprogramming of fibroblasts obtained from a 55-year-old Caucasian female donor. iPSCs were cultured on hESC-qualified Matrigel (Corning) coated vessels in either complete StemFlex (Thermo-Fisher Scientific) or mTeSR Plus (StemCell Technologies). Medium was replaced every other day and cells were passaged ~1:10 onto fresh Matrigel-coated vessels using ReLeSR (StemCell Technologies) when 90% confluent.

### Differentiation of microglia-like cells

Microglia-like cells were differentiated as described (20, 28). Confluent iPSCs were dissociated to single cells using TrypLE (ThermoFisher Scientific). 9 x 10^6^ cells were seeded into 6-well Aggrewell 800 plates (StemCell Technologies) in embryoid body media consisting of complete StemFlex supplemented with 10 μM Y27632 (Stemcell Technologies), 50 ng/mL BMP-4, 50 ng/mL VEGF-121, and 20 ng/mL stem cell factor (SCF) (all from Peprotech). 50% of the media was replaced every other day. Following 7 days in embryoid body media, spheres were lifted from the Aggrewell plate, supernatant was passed through a 40 μm cell strainer (PluriSelect) to remove debris, and the spheres were collected by inverting the strainer into a new conical tube and rinsing with hematopoietic cell media consisting of X-VIVO 15 (Lonza) supplemented with 1X Pen/Strep (Thermo-Fisher Scientific), 1X Glutamax (Thermo-Fisher Scientific), 55 μM β-mercaptoethanol (Gibco), 100 ng/mL M-CSF (Peprotech), and 25 ng/mL IL-3 (Cell Guidance Systems). ~150-300 spheres were seeded into tissue culture-treated T175 flasks in complete hematopoietic cell media. 50% of the media was replaced on a weekly basis. Following ~4 weeks in culture, macrophage precursor cells in suspension were collected from the supernatant upon media change, passed through a 40 μm cell strainer to remove clumps and debris, centrifuged and plated in microglia differentiation medium consisting of Advanced DMEM/F12 (ThermoFisher Scientific) supplemented with 1X Pen/Strep (Thermo-Fisher Scientific), 1X Glutamax (Thermo-Fisher Scientific), 1X N2 supplement (ThermoFisher Scientific), 100 ng/mL IL-34, and 10 ng/mL GM-CSF (both from Peprotech) at 100,000 cells per cm^2^ onto tissue culture-treated dishes, which was again added after 3 days without removing spent medium. Following 7 days in culture, microglia-like cells were removed from the plate using TrypLE and a cell scraper, centrifuged, and cryopreserved in NutriFreeze D10 cryopreservation solution (Biological Industries).

### Differentiation of induced neurons (iNeurons)

Neurons were differentiated using lentiviral-mediated overexpression of the neuronal transcription factor NGN2, per published protocols (24) and as our team has described previously (29). Briefly, confluent iPSC cultures were dissociated to single cells using TrypLE and transduced with a pair of lentiviruses to express rtTA under the control of a ubiquitous promotor and NGN2 under the control of a TetO promotor (pTet-O-NGN2-2A-Puro) at MOIs between 3-5. Infected cells were seeded onto Geltrex (Thermo-Fisher Scientific) coated tissue culture plates in StemFlex media supplemented with 10 μM Y-27632. The following day, media was replaced with fresh StemFlex without the addition of Y-27632, and iPSCs with integrated lentiviruses were expanded and banked in NutriFreeze D10 solution (Biological Industries).

To induce differentiation, confluent infected iPSC cultures were dissociated to single cells using TrypLE and seeded onto Geltrex-coated culture vessels in StemFlex media supplemented with 10 μM Y-27632 and 2 μg/mL Doxycycline (Thermo-Fisher Scientific) to induce TetO-controlled NGN2 expression. 24 hours later media was changed to N2 media consisting of DMEM/F12 supplemented with 1X N2 supplement (Thermo-Fisher Scientific), 1X Glutamax (Thermo-Fisher Scientific), 3% D-(+)-Glucose (Sigma Aldrich), 2 μg/mL Doxycycline, and 3.34 μg/mL Puromycin (Thermo-Fisher Scientific) to begin selection. 48 hours following induction media was changed to N2/B27 media consisting of DMEM/F12 supplemented with 1X N2 supplement, 1X B27 supplement (Thermo-Fisher Scientific), 1X Glutamax, 3% D-(+)-Glucose, 2 μg/mL Doxycycline, and 3.34 μg/mL Puromycin. 72 hours following induction media was replaced with Neurobasal/B27 media consisting of Neurobasal (Thermo-Fisher Scientific) supplemented with 1X B27 supplement, 1X Glutamax, 0.5X non-essential amino acids (Thermo-Fisher Scientific), 3% D-(+)-Glucose, 2 μg/mL Doxycycline, 3.34 μg/mL Puromycin, and 50 ng/ml each of brain-derived neurotrophic factor (BDNF) and glial-derived neurotrophic factor (GDNF) (both from Peprotech). iNeurons were cultured Neurobasal/B27 media for 72 hours without media replacement. 6 days following induction cells were dissociated with Accumax (Millipore), collected and centrifuged, and cryopreserved in Nutrifreeze D10 solution.

### Establishment of neuron-microglia co-cultures

Co-cultures of iNeurons and microglia were established in a customized media amenable to neuron and microglia survival and function consisting of Neurobasal supplemented with 1X nonessential amino acids, 1X Glutamax, 1X N2 supplement, 1X Pen/strep, BDNF (10 ng/mL), GDNF (10 ng/mL), IL-34 (100 ng/mL) and GM-CSF (10 ng/mL) (all protein growth factors were from Peprotech). Prior to thaw, 96-well PDL-coated assay plates (Corning) were coated with mouse laminin at 10 μg/mL and placed in an incubator for ~1 hour. iNeurons and microglia were thawed and seeded at the same time into the laminin coated plate in media at 60,000 neurons and 12,000 microglia per well. 50% of the media was replaced 48 hours after seeding. When indicated, 1 μM final concentration of an LNA GapmeR antisense oligonucleotide (ASO) against *SNCA* (sequence: G*G*C*T*A*A*T*G*A*A*T*T*C*C*T*T), or non-targeting (NT) control (Cat#339515) (Qiagen), were added to the co-culture during the media replacement. Cultures were treated with *SNCA* or NT ASO for 10 days prior to analysis (12 days post-thaw).

### Immunocytochemistry

Immunocytochemistry was performed on monocultures and co-cultures 10-12 days post-thaw. Cells were fixed in 4% paraformaldehyde for 15 minutes at room temperature, washed three times with PBS, and blocked and permeabilized in 10% donkey serum (Abcam) supplemented with 0.3% Triton X-100 (Sigma-Aldrich) for 15 minutes at room temperature. Blocking and permeabilization solution was aspirated and cells were washed once with PBS. Primary antibodies were resuspended in antibody dilution buffer consisting of PBS supplemented with 2% donkey serum, 1% bovine serum albumin (Sigma-Aldrich), 0.2% Tween-20 (Sigma-Aldrich) and incubated overnight at 4°C. The following primary antibodies were used: goat anti-IBA1 (Abcam 5076, 1:500), rabbit anti-CD11b (Abcam ab133357, 1:500), rabbit anti-α-synuclein (Abcam ab138501, 1:150), mouse anti-HuC/D (Thermo-Fisher Scientific A-21271, 1:100), and chicken anti-TUJ1 (Aves TUJ, 1:500). Cells were washed three times with PBS and then visualized with appropriate Alexa-Fluor conjugated secondary antibodies (all from Thermo-Fisher Scientific): Donkey anti-rabbit 555, donkey anti-goat 555, donkey anti-rabbit 647, donkey anti-mouse 555, and donkey anti-chicken 488 (all diluted 1:1000 in antibody dilution buffer). Hoechst (1:2500; Thermo-Fisher Scientific), and where applicable Alexa Fluor 750-conjugated Phalloidin (1:1000, Thermo-Fisher Scientific), were added to the secondary antibody cocktail for visualization of nuclei and F-actin filaments. Secondary antibodies were applied at room temperature for one hour. Cells were washed three times with PBS and imaged on a Cell Insight CX5 or CX7 high content imaging platform (Thermo-Fisher Scientific) in widefield mode using a 10X objective. Quantification of cell populations and α-synuclein fluorescence intensity was performed using Columbus (Perkin Elmer) software by segmenting on viable nuclei and quantifying the fluorescence intensity of the appropriate antibody channel.

### Measurement of SNCA gene expression by qPCR

Gene expression analysis was performed on monocultures and co-cultures 10-12 days post-thaw. Total RNA was extracted using a 1 step Cells-to-CT kit (Thermo-Fisher Scientific) per the manufacturer’s instructions. qRT-PCR was performed on total lysate using a FAM labeled *SNCA* Taqman probe (Thermo-Fisher Scientific, Hs00240906_m1). Expression was normalized to VIC labeled housekeeping gene *POLR2A* (Thermo-Fisher Scientific, Hs00172187_m1). Expression was quantified using the ddCT method and is displayed as fold-change relative to the WT cell condition. qPCR was performed on an Applied Biosystems Quantstudio 7 Pro with the following conditions: 1 cycle of 50°C for 5 minutes, 1 cycle of 95°C for 20 seconds, followed by 40 cycles of 95°C for 3 seconds and 60°C for 30 seconds.

### Phagocytosis of pHrodo conjugated E. coli

Phagocytosis was examined following published protocols (15, 30). pHrodo red labeled *E. coli* (Thermo-Fisher Scientific) was resuspended in PBS to 1 mg/mL, vortexed, and sonicated for five minutes prior to use. pHrodo red *E.coli* was added to each well to at a final concentration of 20 μg/mL. Cells were live imaged on an Incucyte S3 every hour for the indicated times using a 10X objective. Images were acquired from 4-8 wells of a 96 well plate per condition, with 2 or 4 fields per well imaged. For analysis, red objects were segmented, and the mean red fluorescence intensity of the red objects was quantified using Incucyte analysis software. Phagocytosis was assessed 10-12 days following establishment of co-cultures.

### Statistical analysis

Statistical analysis was performed using GraphPad Prism v. 8.3 for Windows (GraphPad Software, San Diego, California USA, www.graphpad.com). Analysis of α-synuclein expression and phagocytosis measured in microglia in monoculture was performed with an unpaired student’s t-test comparing WT and *SNCA* donor-derived cells. Statistical analysis of phagocytosis quantified via live cell imaging was performed with a two-way ANOVA with time and donor genotype (WT or *SNCA*) set as factors, and a post-hoc analysis was performed comparing individual timepoints between cell lines and NT vs. ASO treatment conditions using a Sidak’s multiple comparisons test, per a recommended protocol (31). Statistical significance was set at p < 0.05.

## Results

### WT and SNCA iPSCs efficiently differentiate into microglia and iNeurons

Immunocytochemistry revealed that iPSC-derived microglia-like cells from the WT and *SNCA* triplication donor expressed markers CD11b, IBA1, CD44, and F-actin (Figure 1A). iNeurons generated from the same donor iPSCs expressed neuronal markers HuC/D and Tuj1 (Figure 1B). Quantification of nuclei revealed that over 80% of cells were positive for neuronal marker HuC/D and macrophage markers IBA1 and CD11b (Figure 1C-D). Co-cultures of donor-matched neurons and microglia were assembled and distinct cell populations could be visually identified using antibodies against neuronal makers HuC/D and Tuj1 and macrophage marker CD11b (Figure 1E and inset). Phagocytic competence was confirmed by visualizing pHrodo red fluorescent signal in microglia-like cells within the co-culture via live cell imaging over a 24-hour time course (Figure 1F).

**Figure 1.**
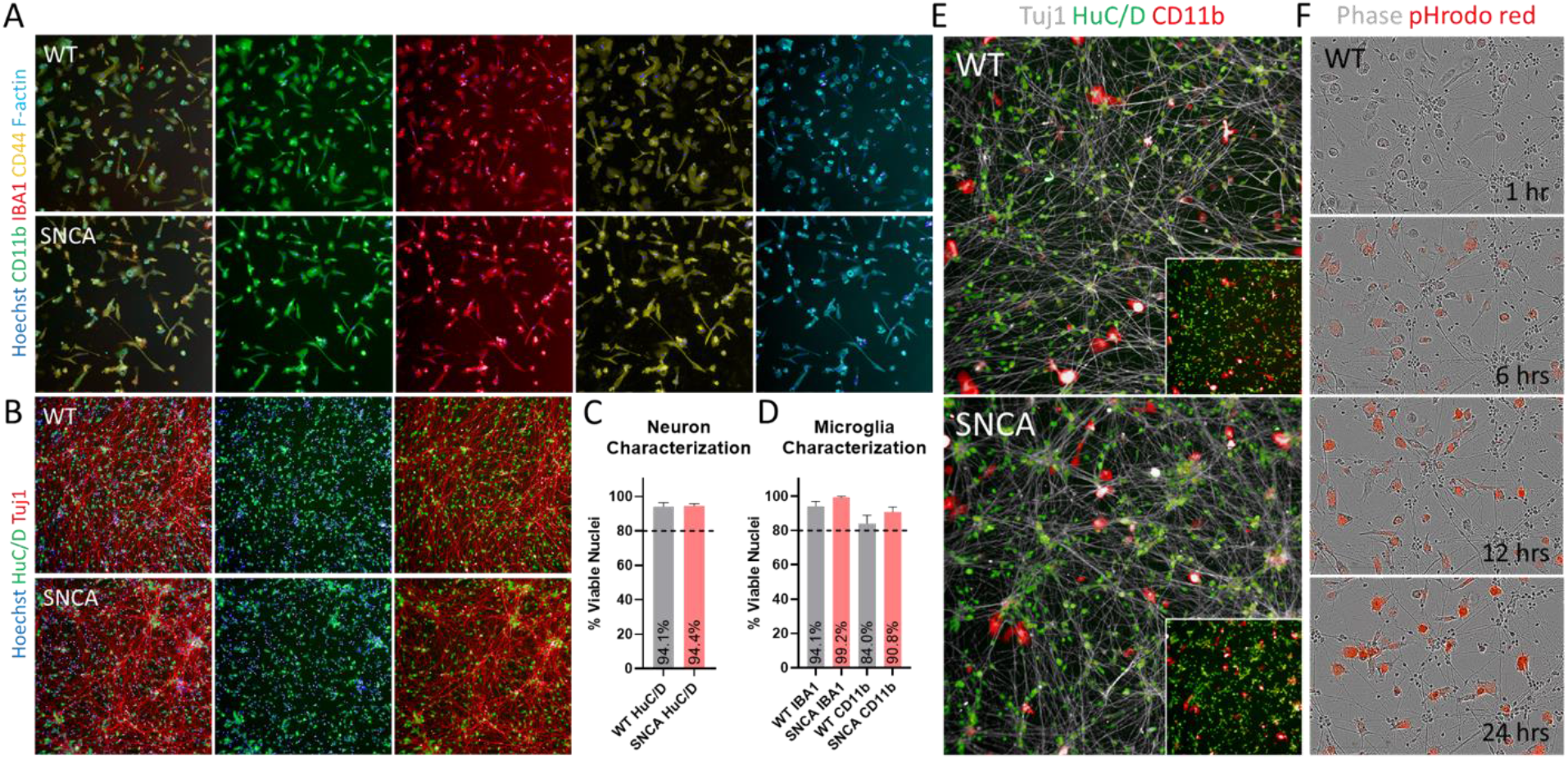
Characterization of microglia-like cells and iNeurons derived from healthy control and SNCA triplication iPSCs. (A-D) Microglia-like cells derived from WT and *SNCA* triplication iPSCs expressed CD11b (green), IBA1 (red), CD44 (yellow), and F-actin (teal). iNeurons derived from WT and *SNCA* triplication iPSCs expressed HuC/D (green) and Tuj1 (red). Greater than 90% of nuclei were positive for the neuronal marker HuC/D, and over 80% of nuclei were positive for microglia markers IBA1 and CD11b. (E) Specific cell types could be visualized in co-cultures of iNeurons and microglia using Tuj1 (grey) and HuC/D (green) to label iNeurons, and CD11b (red) to label microglia. Insets depict distinct HuC/D and CD11b signal in these populations. (F) pHrodo red-labeled E. coli was applied to live co-cultures and imaged over 24-hours. Red signal indicated internalization and localization of the probe to acidic compartments within microglia.

### α-synuclein is elevated in microglia and iNeurons derived from SNCA triplication donor iPSCs

Expression of α-synuclein protein and *SNCA* mRNA was characterized in iPSC-derived microglia and neurons in mono- and co-culture. Fluorescence intensity of α-synuclein protein was significantly increased in *SNCA* triplication donor-derived microglia compared to WT microglia (p < 0.01). *SNCA* transcript level was also ~2.5 fold higher in *SNCA* triplication donor microglia compared to WT microglia (Figure 2A-B). Fluorescence intensity of α-synuclein protein was significantly increased in *SNCA* triplication iNeurons compared to WT iNeurons (p < 0.01). *SNCA* mRNA expression was ~5 fold higher in iNeurons derived from the SNCA donor compared to the WT donor (Figure 2C-D).

**Figure 2.**
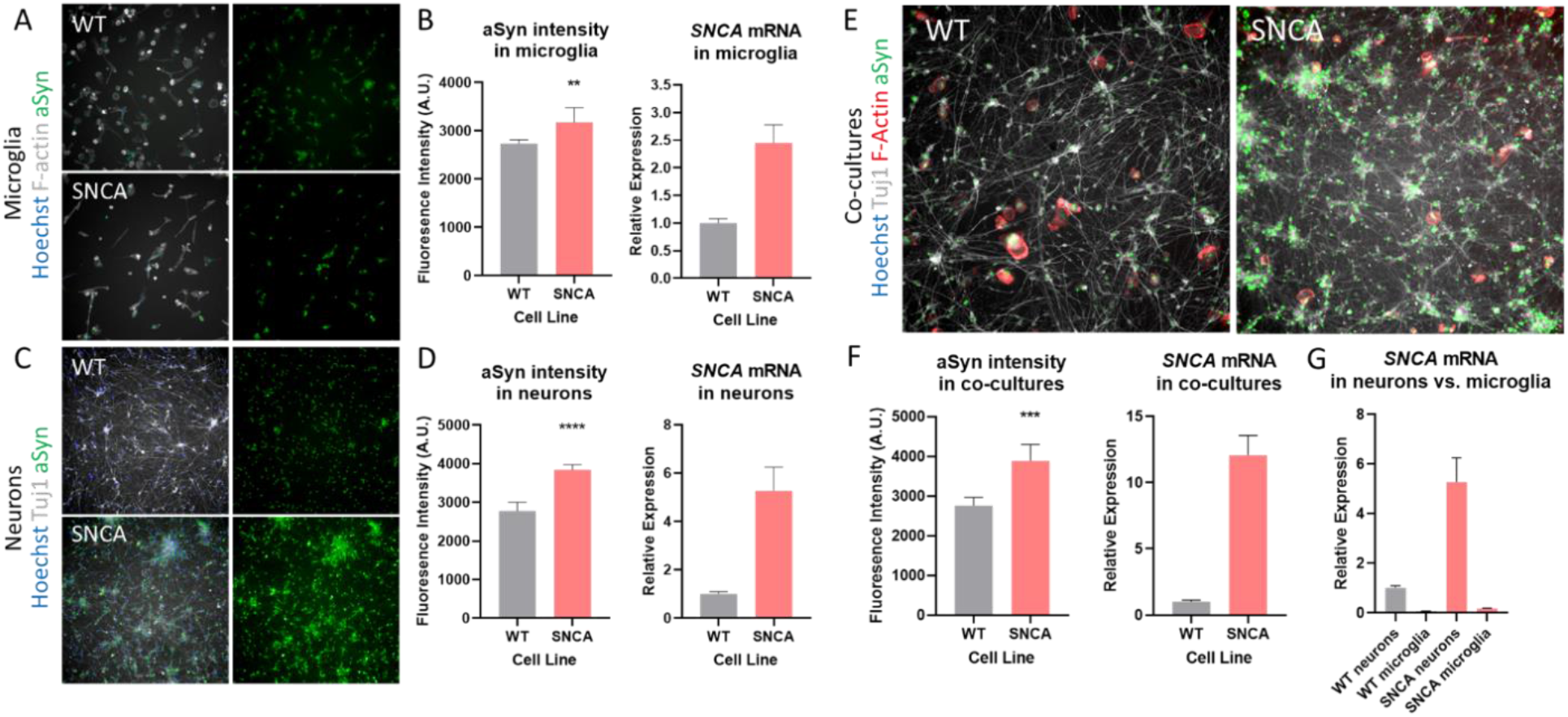
Assessment of α-synuclein expression in microglia and iNeurons derived from healthy control and *SNCA* triplication donor iPSCs. (A) Immunocytochemistry depicting microglia staining for F-actin (grey) and α-synuclein (green). Images to the right depict only α-synuclein signal to highlight difference between donor lines. (B) α-synuclein fluorescence intensity was slightly, but significantly elevated in *SNCA* triplication microglia compared to WT. *SNCA* transcript was ~2.5-fold higher in *SNCA* triplication microglia compared to WT via qPCR. (C) Immunocytochemistry depicting Tuj1 (grey) and α-synuclein (green) in WT and *SNCA* triplication iNeurons. (D) α-synuclein fluorescence intensity was significantly higher in *SNCA* triplication iNeurons compared to WT. qPCR revealed ~5-fold higher *SNCA* transcript expression in *SNCA* triplication iNeurons. (E) Immunocytochemistry depicting Tuj1 (grey) iNeurons, F-actin (red) microglia, and α-synuclein (green) in donor-matched co-cultures. (F) α-synuclein intensity was significantly higher in *SNCA* triplication co-cultures compared to WT. *SNCA* transcript was elevated by ~12-fold in *SNCA* triplication co-cultures compared to WT. (G) *SNCA* transcript was robustly higher in iNeurons compared to microglia in monoculture regardless of donor genotype. Expression is depicted relative to WT iNeurons. * p < 0.05, ** p < 0.01, *** p < 0.001, **** p < 0.0001. Error bars depict ± SD.

Establishment of donor-matched co-cultures did not disrupt this expected phenotype. Intensity of α-synuclein fluorescence was higher in *SNCA* donor-derived co-cultures (p < 0.01) and *SNCA* mRNA was ~12 fold higher in *SNCA* triplication co-cultures compared to WT co-cultures (Figure 2E-F). Assessment of mRNA expression in monocultures across cell types revealed that *SNCA* expression was substantially higher in iNeurons compared to microglia, irrespective of donor genotype (Figure 2G). WT donor iNeurons expressed ~14-fold higher *SNCA* transcript compared to WT donor microglia, while *SNCA* donor iNeurons expressed ~31-fold higher *SNCA* transcript compared to *SNCA* donor microglia.

### Elevated neuronal α-synuclein expression impacts microglia phagocytosis of pHrodo red-labeled E. coli

iPSC-derived microglia phagocytic function was assessed via live cell imaging of pHrodo red-labeled *E. coli*, which only fluoresces when localized to intracellular acidic environments. To assess effects of excess neuronal *SNCA* expression, microglial phagocytosis was examined over 48-hours in co-cultures constructed of donor-matched WT and *SNCA* triplication microglia and iNeurons and mixed co-cultures consisting of *SNCA* triplication iNeurons and WT microglia (Figure 3A). A two-way ANOVA revealed an overall significant effect of time (p < 0.0001) and genotype (p < 0.0001), and an interaction between time and genotype (p <0.0001). Post-hoc statistical analysis comparing donor-matched WT and *SNCA* triplication co-cultures highlighted significantly lower pHrodo red signal in *SNCA* triplication co-cultures (Figure 3B, and Supplemental Table 1). Similarly, post-hoc analysis comparing donor-matched WT co-cultures to mixed co-cultures containing WT microglia and *SNCA* triplication iNeurons highlighted significantly lower pHrodo red signal in mixed co-cultures (Figure 3B, and Supplemental Table 1).

**Figure 3.**
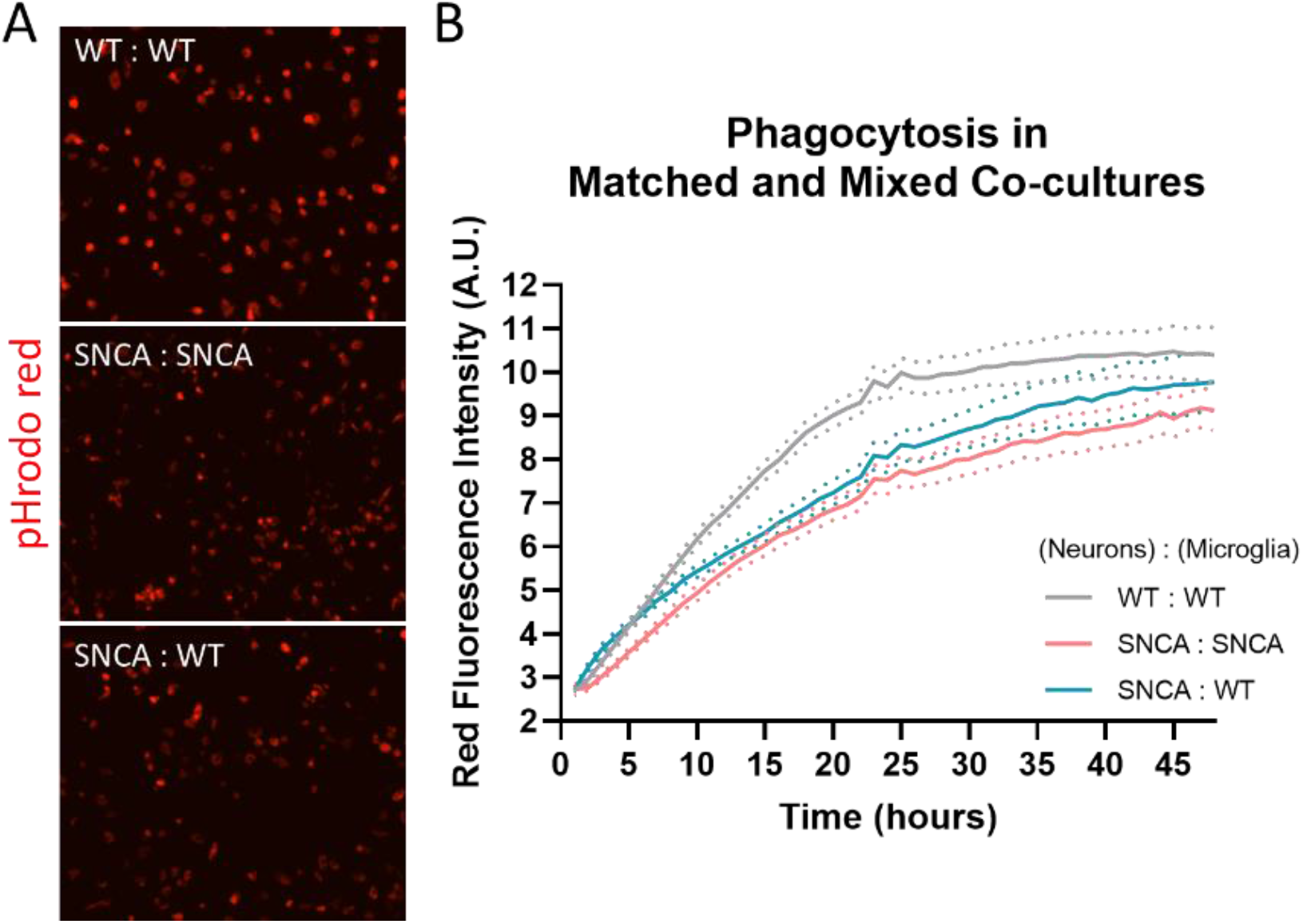
Phagocytosis of pHrodo red *E. coli* is impaired in co-cultures containing *SNCA* triplication donor iNeurons. (A) Representative images after 48-hour incubation of pHrodo red E. coli in co-cultures of neurons:microglia. *Top: WT donor-matched; Middle: SNCA triplication donor-matched; Bottom: SNCA triplication iNeurons co-cultured with WT microglia*. (B) Significantly lower pHrodo red signal was measured in donor matched co-cultures of iNeurons and microglia derived from the *SNCA* triplication donor compared to the WT donor, as well as in mixed co-cultures containing *SNCA* triplication iNeurons and WT microglia compared to WT:WT matched co-cultures. Post-hoc statistics comparing each time point can be found in Supplemental Table 1. Error bars depict ± SD.

### Genetic knockdown of SNCA augments phagocytosis of pHrodo red-labeled E. coli in SNCA triplication co-cultures

Knockdown of *SNCA* mRNA in neurons, microglia, and co-cultures was achieved using an antisense oligonucleotide (ASO) in *SNCA* triplication cell types. Compared to a non-targeting control, we observed a 72% reduction of mRNA in iNeurons, an 81% reduction of mRNA in microglia, and a 57% reduction of mRNA in co-cultures after 10 days of treatment with an *SNCA* ASO (Figure 4A). Immunocytochemistry demonstrated a reduction of α-synuclein protein expression via fluorescence intensity in Tuj1+ iNeurons following *SNCA* ASO treatment in co-cultures (Figure 4B).

**Figure 4.**
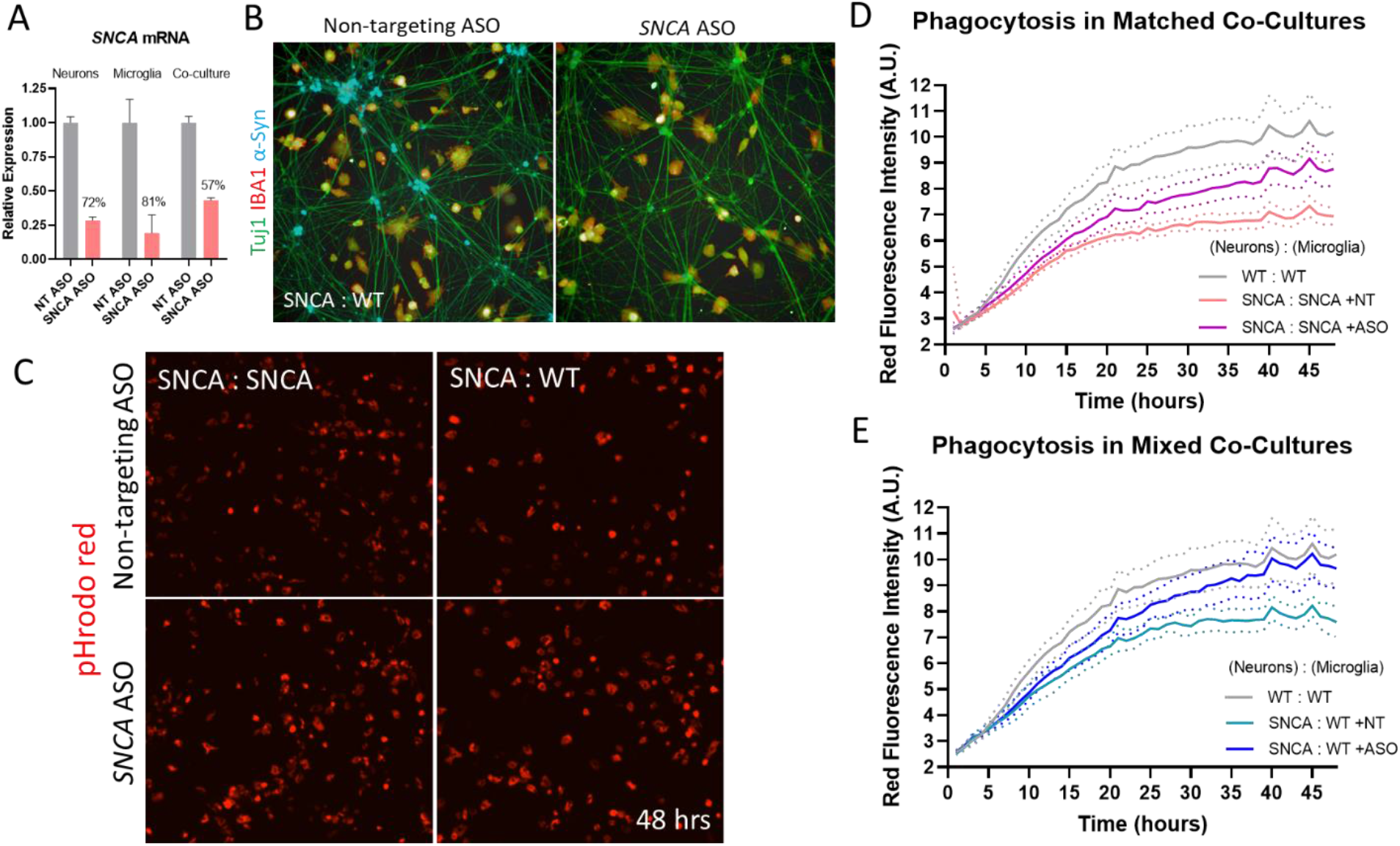
Genetic knockdown of *SNCA* enhances phagocytosis of pHrodo red E. coli in microglia cultured with *SNCA* triplication iNeurons. (A) An antisense oligonucleotide (ASO) against *SNCA* substantially reduced expression of *SNCA* mRNA in iNeurons, microglia, and co-culture in *SNCA* triplication donor cell types. The percent reduction is indicated above the bar. Expression is depicted as relative the non-targeting (NT) control for each cell culture condition. (B) Representative image showing reduced α-synuclein (teal) in co-cultures of iNeurons and microglia. α-synuclein could be visualized in the soma of Tuj1+ iNeurons, and the signal was visibly reduced following 10-day *SNCA* ASO treatment. (C) Representative images after 48-hour incubation of pHrodo red *E. coli* in co-cultures of *SNCA* triplication iNeurons and either *SNCA* triplication or WT microglia. *Top row: non-targeting ASO treated. Bottom row: SNCA ASO treated*. (D-E) In agreement with our previous result, donor-matched or mixed co-cultures containing *SNCA* triplication iNeurons demonstrated significantly attenuated pHrodo red signal compared to WT co-cultures. 10-day treatment with an ASO against *SNCA* significantly increased the fluorescence intensity of pHrodo red signal compared to non-targeting control in co-cultures containing *SNCA* triplication iNeurons and donor matched microglia (D) or mixed co-cultures of *SNCA* triplication iNeurons and WT microglia (E). Post-hoc statistics comparing each time point can be found in Supplemental Tables 2 and 3. Error bars depict ± SD.

The effect of *SNCA* gene expression reduction on microglial function was examined using the pHrodo red phagocytosis assay. Knockdown of *SNCA* with the ASO resulted in a significant increase in pHrodo red intensity in donor matched and mixed co-cultures containing *SNCA* triplication iNeurons (Figure 4C). In donor matched *SNCA* co-cultures, two-way ANOVA revealed a significant effect of time (p < 0.0001) and genotype (p < 0.0001), and an interaction between time and genotype (p <0.0001). Post-hoc analysis revealed that, compared to WT co-cultures, *SNCA* triplication co-cultures again had significantly lower pHrodo red signal (Figure 4D, Supplemental Table 2) and that *SNCA* knockdown resulted in significantly higher pHrodo red signal compared to non-targeting control (Figure 4D, Supplemental Table 2), although we note that intensity remained statistically lower compared to WT co-cultures. In mixed co-cultures containing *SNCA* triplication iNeurons and WT microglia, two-way ANOVA revealed a significant effect of time (p < 0.0001) and genotype (p < 0.0001), and an interaction between time and genotype (p <0.0001). Compared to the WT donor-matched co-cultures, post-hoc analysis again suggested that *SNCA* triplication iNeuron containing co-cultures had significantly lower pHrodo red signal (Figure 4E, Supplemental Table 3) and that *SNCA* knockdown in the mixed co-cultures resulted in significantly higher pHrodo red signal compared to non-targeting control (Figure 4E, Supplemental Table 3). We note that at the later timepoints examined there was not a statistical difference in the ASO-treated mixed co-cultures compared to the WT donor-matched co-cultures.

## Discussion

While recent reports have implicated microglia in the pathogenesis of multiple neurodegenerative diseases, the specific mechanisms underlying their impact on the progression of synucleinopathies remains incompletely understood (13, 32, 33). Novel, disease-relevant model systems that recapitulate the pathologic condition might help to elucidate the mechanisms underlying their contribution. In the current study, we utilized human iPSCs and established differentiation methodologies (20, 24) to generate both microglia-like cells and iNeurons from a healthy control donor and a donor with an allelic triplication of the *SNCA* gene locus, which results in twice the normal amount of the α-synuclein-encoding gene, *SNCA*, and is associated with development of synucleinopathies including Parkinson’s disease (34). iPSCs from both WT and *SNCA* triplication donors were able to efficiently differentiate into cultures of highly pure microglia and iNeurons, which recapitulated the expected phenotype of increased *SNCA* transcript and α-synuclein protein expression in the patient-derived cells. Co-cultures containing donor-matched and mixed cell types (*SNCA* triplication iNeurons and WT microglia) were constructed and microglia functionality was validated using a pHrodo red live cell imaging assay that has been previously described (30). Our results confirmed that we were able to efficiently generate patient-derived cell types that could be used to investigate the effect of excess *SNCA* expression on microglia phagocytosis in a disease-relevant context.

Complementary to the findings of reports suggesting that increased α-synuclein, either due to exogenous application/overexpression or *SNCA* gene triplication, compromises phagocytosis in human iPSC-derived macrophages (15) and immortalized murine BV2 microglia (16), we observed reduced phagocytosis of pHrodo red *E. coli* in co-cultures constructed of *SNCA* triplication donor-derived neurons and microglia compared to co-cultures constructed from the WT donor. A novel finding of our study was that we were able to phenocopy the attenuated phagocytosis of *SNCA* triplication co-cultures when genetically normal WT microglia were co-cultured with the *SNCA* triplication iNeurons. This finding is particularly interesting in the context of our result that *SNCA* gene expression is substantially higher in iNeurons compared to microglia, regardless of donor genotype (Figure 2G), and suggests that excess neuronal α-synuclein, either directly or through undetermined downstream consequences, is driving the phenotype observed in microglia via non-cell autonomous mechanisms in our *in vitro* model system. Importantly, we highlight that we observed this phenotype in our iPSC co-culture model without the supraphysiologic overexpression or exogenous application of α-synuclein, but instead through construction of disease-relevant cell types that capture the genetics of human disease. Our findings are in agreement with the discussions of Booms and Coertzee (2021) in that monitoring microglia function under disease- and genetically-relevant physiological conditions seen in humans is important to better understand and elucidate how endogenous *SNCA* influences microglia function in PD (35).

Human genetics data indicate an *SNCA* gene dosage-effect on PD pathology, with patients harboring a triplication of the *SNCA* locus having a more severe phenotype and earlier disease onset than patients harboring a duplication of this region (3, 6–8). In support of these seminal observations, human and animal studies have suggested that downregulation of the *SNCA* gene is therapeutically efficacious in modifying pathology in the disease setting (36–39). To test our hypothesis that excess neuronal α-synuclein was driving the observed phagocytosis phenotype in our co-culture system, we knocked down *SNCA* gene expression using an antisense oligonucleotide (ASO). We were able to achieve ~50% knockdown of *SNCA* in co-cultures, which is in line with *in vivo* models that demonstrate efficacy and within a proposed therapeutic range (37, 40, 41). We observed that knockdown of *SNCA* resulted in higher pHrodo red signal in *SNCA* triplication donor-matched co-cultures. Importantly, we also observed that knockdown of *SNCA* augmented phagocytosis in our mixed co-cultures constructed of WT microglia and *SNCA* triplication iNeurons that phenocopied the *SNCA* triplication donor matched cultures. These results suggest that the attenuated phagocytic ability of microglia cultured with *SNCA* triplication iNeurons can be manipulated by modifying the expression of *SNCA* transcript within the culture. A potential future use of this assay could be to further define the relationship between *SNCA* expression level and phagocytic competence by applying a range of concentrations of *SNCA* ASOs to modulate gene expression to varying degrees.

Taken all together and in the context of the prior publications cited above, our results suggest that excess levels of neuronal-derived α-synuclein generated because of a disease-relevant *SNCA* locus triplication impacts phagocytic function of microglia in or model system. This conclusion supports the work of Haenseler et al. (2017), which demonstrated that *SNCA* triplication, but not an *SNCA* A53T point mutation that does not lead to excess synuclein transcription, resulted in impaired phagocytosis in iPSC-derived macrophages (15). We acknowledge a limitation of our data in that we cannot infer whether direct contact between neurons and microglia is required to observe this phenotype, or what, if any, secreted neuronal factors are impacting microglia function. While we speculate that excess secreted α-synuclein protein derived from the *SNCA* triplication iNeurons is a candidate, we cannot rule out that other factors downstream of elevated α-synuclein, such as inflammatory cytokines produced as a response to exposure (42), might be mediating this effect. Future experiments could be designed to address this question by examining the effects of conditioned media harvested from *SNCA* triplication iNeurons and applying it to microglia in monoculture prior to measuring phagocytosis, or by directly interrogating the components of the conditioned media via proteomics.

Phagocytic competence of microglia is critical for maintaining CNS homeostasis, and perturbation of this process is a hallmark observed across many disease states (43). Indeed, impairment of microglial phagocytosis has been observed in human samples and PD models (for review, see (44)) and postulated as a modifiable therapeutic strategy (13). Some key pieces of evidence in support of this notion come from examining variants in *TREM2*, which encodes a transmembrane receptor expressed on microglia, among other cells, that upon activation signals to modulate microglial function and inflammatory state (45). Missense mutations in *TREM2* impair phagocytosis in iPSC-derived microglia (46) and have been associated with the development of PD (47). In *in vitro* and *in vivo* models of PD, *TREM2* insufficiency augmented inflammatory responses and apoptosis in cell models, and exacerbated *SNCA* overexpression-induced neuronal loss and shifted microglia to a proinflammatory state in rodents (48). While the effects of *TREM2* variants can impact microglial processes in addition to phagocytosis, augmenting phagocytosis with a PPARγ agonist was shown to improve both cellular and behavioral outcomes in an MPTP-induced mouse model of PD, suggesting that maintaining normal phagocytic competence could be beneficial in PD (49). Taken all together, the role and importance of microglia and phagocytic processes in PD progression remain incompletely understood. We propose that our novel, patient-derived co-culture system can be a useful tool in both further understanding the underlying relationships between aberrant *SNCA* expression and microglia phagocytosis, and in identifying or validating therapeutic agents that aim to modify the expression of *SNCA* or enhance the phagocytic ability of microglia in a disease-relevant cellular and genetic context.

The results of our studies should be interpreted with respect to its limitations. First, NGN2 overexpression to generate iNeurons from iPSCs produces heterogeneous mixed cultures enriched for excitatory neurons (50), which are not necessarily the most relevant neuronal subtype associated with the onset and progression of Parkinson’s disease. Establishment of co-cultures containing donor matched microglia and dopaminergic neurons would perhaps better model the disease state. Second, we chose to examine phagocytosis using an assay established in prior studies that took advantage of the commercial availably of pHrodo conjugated *E. coli*, which is not a disease-relevant substrate. Prior reports have in fact noted discrepancies in phagocytosis of different substrates across iPSC-derived microglia generated from donors with TREM2 mutations (46). Future studies could harness these novel co-culture systems to examine phagocytosis using substrates relevant to the PD state, such as neuronal debris or fluorescent α-synuclein. Third, we used WT and *SNCA* triplication iPSCs that were derived from separate donors, and therefore the differentiate progeny might inherently have differences in function that are not associated with disease mutations. We sought to overcome this limitation by using an ASO to reduce *SNCA* expression in our *SNCA* triplication disease model to more physiologic level (~57% reduction). However, co-cultures constructed from differentiated cells derived from isogenic iPSC lines that have been edited to contain the disease mutation would be a gold-standard model system that can be utilized in subsequent studies. Finally, our investigation does not identify the biological mechanisms that leads to impaired microglial phagocytosis in the presence of *SNCA* triplication iNeurons. We suggest that utilization of the co-culture system described here, or perhaps examination of phagocytosis in the presence of *SNCA* iNeuron conditioned media, could help to elucidate an underlying molecular cascade evoked by α-synuclein that results in this phenotype, and perhaps a better understanding of how excess α-synuclein might perturb microglial function.

In summary, we have leveraged iPSC technologies to generate both iNeurons and microglia from a healthy control donor and a Parkinson’s disease donor with an allelic triplication of the *SNCA* gene locus. With these tools, we constructed co-culture models that recapitulated the disease state, and found that microglial phagocytosis was impaired in co-cultures containing *SNCA* triplication iNeurons. Reducing *SNCA* expression partially ameliorated this phenotype, suggesting a detrimental role of excess neuronal *SNCA* on microglial function. The results of our studies complement the previous findings and conclusions in the field, but importantly extend upon these previous works by describing this phenotype in a physiologically- and genetically-relevant co-culture model system.

---

**Supplemental Table 1:** Sidak’s Multiple Comparisons Post-hoc Test of Phagocytosis in Matched and Mixed Co-Cultures.

|  |  |  |  |  |  |
| --- | --- | --- | --- | --- | --- |
| Sidak's multiple comparisons test | Mean Diff. | 95.00% CI of diff. | Below threshold? | Summary | Adjusted P Value |
| Hour 1 |  |  |  |  |  |
| WT : WT vs. SNCA : SNCA | -0.02413 | -0.4435 to 0.3952 | No | ns | 0.9895 |
| WT : WT vs. SNCA : WT | -0.02609 | -0.4455 to 0.3933 | No | ns | 0.9877 |
| Hour 2 |  |  |  |  |  |
| WT : WT vs. SNCA : SNCA | 0.1975 | -0.2219 to 0.6168 | No | ns | 0.4984 |
| WT : WT vs. SNCA : WT | -0.2727 | -0.6921 to 0.1466 | No | ns | 0.2698 |
| Hour 3 |  |  |  |  |  |
| WT : WT vs. SNCA : SNCA | 0.3317 | -0.08770 to 0.7510 | No | ns | 0.1477 |
| WT : WT vs. SNCA : WT | -0.2911 | -0.7104 to 0.1283 | No | ns | 0.2262 |
| Hour 4 |  |  |  |  |  |
| WT : WT vs. SNCA : SNCA | 0.4885 | 0.06911 to 0.9078 | Yes | * | 0.0183 |
| WT : WT vs. SNCA : WT | -0.1623 | -0.5817 to 0.2570 | No | ns | 0.6232 |
| Hour 5 |  |  |  |  |  |
| WT : WT vs. SNCA : SNCA | 0.6043 | 0.1849 to 1.024 | Yes | ** | 0.0026 |
| WT : WT vs. SNCA : WT | -0.03181 | -0.4512 to 0.3875 | No | ns | 0.9818 |
| Hour 6 |  |  |  |  |  |
| WT : WT vs. SNCA : SNCA | 0.7218 | 0.3024 to 1.141 | Yes | *** | 0.0002 |
| WT : WT vs. SNCA : WT | 0.1214 | -0.2980 to 0.5408 | No | ns | 0.7666 |
| Hour 7 |  |  |  |  |  |
| WT : WT vs. SNCA : SNCA | 0.8605 | 0.4412 to 1.280 | Yes | **** | <0.0001 |
| WT : WT vs. SNCA : WT | 0.2445 | -0.1749 to 0.6638 | No | ns | 0.3471 |
| Hour 8 |  |  |  |  |  |
| WT : WT vs. SNCA : SNCA | 0.9856 | 0.5662 to 1.405 | Yes | **** | <0.0001 |
| WT : WT vs. SNCA : WT | 0.4375 | 0.01815 to 0.8569 | Yes | * | 0.0389 |
| Hour 9 |  |  |  |  |  |
| WT : WT vs. SNCA : SNCA | 1.08 | 0.6608 to 1.499 | Yes | **** | <0.0001 |
| WT : WT vs. SNCA : WT | 0.5498 | 0.1305 to 0.9692 | Yes | ** | 0.0068 |
| Hour 10 |  |  |  |  |  |
| WT : WT vs. SNCA : SNCA | 1.243 | 0.8232 to 1.662 | Yes | **** | <0.0001 |
| WT : WT vs. SNCA : WT | 0.7418 | 0.3224 to 1.161 | Yes | *** | 0.0002 |
| Hour 11 |  |  |  |  |  |
| WT : WT vs. SNCA : SNCA | 1.308 | 0.8884 to 1.727 | Yes | **** | <0.0001 |
| WT : WT vs. SNCA : WT | 0.8999 | 0.4805 to 1.319 | Yes | **** | <0.0001 |
| Hour 12 |  |  |  |  |  |
| WT : WT vs. SNCA : SNCA | 1.35 | 0.9311 to 1.770 | Yes | **** | <0.0001 |
| WT : WT vs. SNCA : WT | 0.9718 | 0.5524 to 1.391 | Yes | **** | <0.0001 |
| Hour 13 |  |  |  |  |  |
| WT : WT vs. SNCA : SNCA | 1.43 | 1.010 to 1.849 | Yes | **** | <0.0001 |
| WT : WT vs. SNCA : WT | 1.111 | 0.6916 to 1.530 | Yes | **** | <0.0001 |
| Hour 14 |  |  |  |  |  |
| WT : WT vs. SNCA : SNCA | 1.577 | 1.158 to 1.996 | Yes | **** | <0.0001 |
| WT : WT vs. SNCA : WT | 1.274 | 0.8546 to 1.693 | Yes | **** | <0.0001 |
| Hour 15 |  |  |  |  |  |
| WT : WT vs. SNCA : SNCA | 1.709 | 1.290 to 2.129 | Yes | **** | <0.0001 |
| WT : WT vs. SNCA : WT | 1.424 | 1.005 to 1.843 | Yes | **** | <0.0001 |
| Hour 16 |  |  |  |  |  |
| WT : WT vs. SNCA : SNCA | 1.725 | 1.305 to 2.144 | Yes | **** | <0.0001 |
| WT : WT vs. SNCA : WT | 1.438 | 1.019 to 1.858 | Yes | **** | <0.0001 |
| Hour 17 |  |  |  |  |  |
| WT : WT vs. SNCA : SNCA | 1.939 | 1.520 to 2.359 | Yes | **** | <0.0001 |
| WT : WT vs. SNCA : WT | 1.605 | 1.185 to 2.024 | Yes | **** | <0.0001 |
| Hour 18 |  |  |  |  |  |
| WT : WT vs. SNCA : SNCA | 2.092 | 1.673 to 2.511 | Yes | **** | <0.0001 |
| WT : WT vs. SNCA : WT | 1.738 | 1.318 to 2.157 | Yes | **** | <0.0001 |
| Hour 19 |  |  |  |  |  |
| WT : WT vs. SNCA : SNCA | 2.114 | 1.695 to 2.534 | Yes | **** | <0.0001 |
| WT : WT vs. SNCA : WT | 1.727 | 1.308 to 2.146 | Yes | **** | <0.0001 |
| Hour 20 |  |  |  |  |  |
| WT : WT vs. SNCA : SNCA | 2.163 | 1.744 to 2.582 | Yes | **** | <0.0001 |
| WT : WT vs. SNCA : WT | 1.78 | 1.360 to 2.199 | Yes | **** | <0.0001 |
| Hour 21 |  |  |  |  |  |
| WT : WT vs. SNCA : SNCA | 2.207 | 1.788 to 2.627 | Yes | **** | <0.0001 |
| WT : WT vs. SNCA : WT | 1.728 | 1.309 to 2.148 | Yes | **** | <0.0001 |
| Hour 22 |  |  |  |  |  |
| WT : WT vs. SNCA : SNCA | 2.154 | 1.735 to 2.574 | Yes | **** | <0.0001 |
| WT : WT vs. SNCA : WT | 1.711 | 1.292 to 2.131 | Yes | **** | <0.0001 |
| Hour 23 |  |  |  |  |  |
| WT : WT vs. SNCA : SNCA | 2.256 | 1.836 to 2.675 | Yes | **** | <0.0001 |
| WT : WT vs. SNCA : WT | 1.708 | 1.289 to 2.127 | Yes | **** | <0.0001 |
| Hour 24 |  |  |  |  |  |
| WT : WT vs. SNCA : SNCA | 2.131 | 1.711 to 2.550 | Yes | **** | <0.0001 |
| WT : WT vs. SNCA : WT | 1.619 | 1.200 to 2.039 | Yes | **** | <0.0001 |
| Hour 25 |  |  |  |  |  |
| WT : WT vs. SNCA : SNCA | 2.246 | 1.827 to 2.666 | Yes | **** | <0.0001 |
| WT : WT vs. SNCA : WT | 1.655 | 1.236 to 2.074 | Yes | **** | <0.0001 |
| Hour 26 |  |  |  |  |  |
| WT : WT vs. SNCA : SNCA | 2.21 | 1.790 to 2.629 | Yes | **** | <0.0001 |
| WT : WT vs. SNCA : WT | 1.582 | 1.163 to 2.002 | Yes | **** | <0.0001 |
| Hour 27 |  |  |  |  |  |
| WT : WT vs. SNCA : SNCA | 2.108 | 1.689 to 2.528 | Yes | **** | <0.0001 |
| WT : WT vs. SNCA : WT | 1.483 | 1.064 to 1.902 | Yes | **** | <0.0001 |
| Hour 28 |  |  |  |  |  |
| WT : WT vs. SNCA : SNCA | 2.109 | 1.690 to 2.529 | Yes | **** | <0.0001 |
| WT : WT vs. SNCA : WT | 1.461 | 1.041 to 1.880 | Yes | **** | <0.0001 |
| Hour 29 |  |  |  |  |  |
| WT : WT vs. SNCA : SNCA | 1.977 | 1.558 to 2.397 | Yes | **** | <0.0001 |
| WT : WT vs. SNCA : WT | 1.368 | 0.9482 to 1.787 | Yes | **** | <0.0001 |
| Hour 30 |  |  |  |  |  |
| WT : WT vs. SNCA : SNCA | 2.021 | 1.601 to 2.440 | Yes | **** | <0.0001 |
| WT : WT vs. SNCA : WT | 1.324 | 0.9042 to 1.743 | Yes | **** | <0.0001 |
| Hour 31 |  |  |  |  |  |
| WT : WT vs. SNCA : SNCA | 1.984 | 1.565 to 2.404 | Yes | **** | <0.0001 |
| WT : WT vs. SNCA : WT | 1.335 | 0.9152 to 1.754 | Yes | **** | <0.0001 |
| Hour 32 |  |  |  |  |  |
| WT : WT vs. SNCA : SNCA | 1.934 | 1.515 to 2.354 | Yes | **** | <0.0001 |
| WT : WT vs. SNCA : WT | 1.209 | 0.7895 to 1.628 | Yes | **** | <0.0001 |
| Hour 33 |  |  |  |  |  |
| WT : WT vs. SNCA : SNCA | 1.845 | 1.426 to 2.265 | Yes | **** | <0.0001 |
| WT : WT vs. SNCA : WT | 1.23 | 0.8111 to 1.650 | Yes | **** | <0.0001 |
| Hour 34 |  |  |  |  |  |
| WT : WT vs. SNCA : SNCA | 1.78 | 1.361 to 2.200 | Yes | **** | <0.0001 |
| WT : WT vs. SNCA : WT | 1.114 | 0.6945 to 1.533 | Yes | **** | <0.0001 |
| Hour 35 |  |  |  |  |  |
| WT : WT vs. SNCA : SNCA | 1.848 | 1.428 to 2.267 | Yes | **** | <0.0001 |
| WT : WT vs. SNCA : WT | 1.04 | 0.6211 to 1.460 | Yes | **** | <0.0001 |
| Hour 36 |  |  |  |  |  |
| WT : WT vs. SNCA : SNCA | 1.776 | 1.356 to 2.195 | Yes | **** | <0.0001 |
| WT : WT vs. SNCA : WT | 1.025 | 0.6053 to 1.444 | Yes | **** | <0.0001 |
| Hour 37 |  |  |  |  |  |
| WT : WT vs. SNCA : SNCA | 1.69 | 1.270 to 2.109 | Yes | **** | <0.0001 |
| WT : WT vs. SNCA : WT | 0.9994 | 0.5801 to 1.419 | Yes | **** | <0.0001 |
| Hour 38 |  |  |  |  |  |
| WT : WT vs. SNCA : SNCA | 1.784 | 1.365 to 2.204 | Yes | **** | <0.0001 |
| WT : WT vs. SNCA : WT | 0.9454 | 0.5261 to 1.365 | Yes | **** | <0.0001 |
| Hour 39 |  |  |  |  |  |
| WT : WT vs. SNCA : SNCA | 1.705 | 1.286 to 2.124 | Yes | **** | <0.0001 |
| WT : WT vs. SNCA : WT | 1.028 | 0.6087 to 1.447 | Yes | **** | <0.0001 |
| Hour 40 |  |  |  |  |  |
| WT : WT vs. SNCA : SNCA | 1.684 | 1.264 to 2.103 | Yes | **** | <0.0001 |
| WT : WT vs. SNCA : WT | 0.8972 | 0.4778 to 1.317 | Yes | **** | <0.0001 |
| Hour 41 |  |  |  |  |  |
| WT : WT vs. SNCA : SNCA | 1.635 | 1.215 to 2.054 | Yes | **** | <0.0001 |
| WT : WT vs. SNCA : WT | 0.8755 | 0.4562 to 1.295 | Yes | **** | <0.0001 |
| Hour 42 |  |  |  |  |  |
| WT : WT vs. SNCA : SNCA | 1.624 | 1.205 to 2.043 | Yes | **** | <0.0001 |
| WT : WT vs. SNCA : WT | 0.7884 | 0.3690 to 1.208 | Yes | **** | <0.0001 |
| Hour 43 |  |  |  |  |  |
| WT : WT vs. SNCA : SNCA | 1.473 | 1.054 to 1.893 | Yes | **** | <0.0001 |
| WT : WT vs. SNCA : WT | 0.782 | 0.3627 to 1.201 | Yes | **** | <0.0001 |
| Hour 44 |  |  |  |  |  |
| WT : WT vs. SNCA : SNCA | 1.36 | 0.9407 to 1.779 | Yes | **** | <0.0001 |
| WT : WT vs. SNCA : WT | 0.7898 | 0.3704 to 1.209 | Yes | **** | <0.0001 |
| Hour 45 |  |  |  |  |  |
| WT : WT vs. SNCA : SNCA | 1.533 | 1.113 to 1.952 | Yes | **** | <0.0001 |
| WT : WT vs. SNCA : WT | 0.7692 | 0.3499 to 1.189 | Yes | **** | <0.0001 |
| Hour 46 |  |  |  |  |  |
| WT : WT vs. SNCA : SNCA | 1.316 | 0.8965 to 1.735 | Yes | **** | <0.0001 |
| WT : WT vs. SNCA : WT | 0.7056 | 0.2862 to 1.125 | Yes | *** | 0.0003 |
| Hour 47 |  |  |  |  |  |
| WT : WT vs. SNCA : SNCA | 1.255 | 0.8355 to 1.674 | Yes | **** | <0.0001 |
| WT : WT vs. SNCA : WT | 0.6833 | 0.2639 to 1.103 | Yes | *** | 0.0006 |
| Hour 48 |  |  |  |  |  |
| WT : WT vs. SNCA : SNCA | 1.26 | 0.8407 to 1.679 | Yes | **** | <0.0001 |
| WT : WT vs. SNCA : WT | 0.6233 | 0.2039 to 1.043 | Yes | ** | 0.0018 |

**Supplemental Table 2:** Sidak’s Multiple Comparisons Post-hoc Test of Phagocytosis Following ASO Treatment in Matched Co-Cultures.

|  |  |  |  |  |  |
| --- | --- | --- | --- | --- | --- |
| Sidak's multiple comparisons test | Mean Diff. | 95.00% CI of diff. | Below threshold? | Summary | Adjusted P Value |
| Hour 1 |  |  |  |  |  |
| WT : WT vs. SNCA : SNCA +NT | -0.7114 | -1.355 to -0.06810 | Yes | * | 0.0247 |
| WT : WT vs. SNCA : SNCA +ASO | -0.04129 | -0.6846 to 0.6020 | No | ns | 0.9982 |
| SNCA : SNCA +NT vs. SNCA : SNCA +ASO | 0.6701 | 0.02681 to 1.313 | Yes | * | 0.0382 |
| Hour 2 |  |  |  |  |  |
| WT : WT vs. SNCA : SNCA +NT | 0.01741 | -0.6259 to 0.6607 | No | ns | 0.9999 |
| WT : WT vs. SNCA : SNCA +ASO | -0.03246 | -0.6758 to 0.6109 | No | ns | 0.9991 |
| SNCA : SNCA +NT vs. SNCA : SNCA +ASO | -0.04987 | -0.6932 to 0.5934 | No | ns | 0.9968 |
| Hour 3 |  |  |  |  |  |
| WT : WT vs. SNCA : SNCA +NT | 0.1013 | -0.5420 to 0.7446 | No | ns | 0.9747 |
| WT : WT vs. SNCA : SNCA +ASO | -0.01228 | -0.6556 to 0.6310 | No | ns | >0.9999 |
| SNCA : SNCA +NT vs. SNCA : SNCA +ASO | -0.1136 | -0.7569 to 0.5297 | No | ns | 0.965 |
| Hour 4 |  |  |  |  |  |
| WT : WT vs. SNCA : SNCA +NT | 0.1298 | -0.5135 to 0.7731 | No | ns | 0.9491 |
| WT : WT vs. SNCA : SNCA +ASO | 0.02425 | -0.6191 to 0.6676 | No | ns | 0.9996 |
| SNCA : SNCA +NT vs. SNCA : SNCA +ASO | -0.1056 | -0.7489 to 0.5377 | No | ns | 0.9716 |
| Hour 5 |  |  |  |  |  |
| WT : WT vs. SNCA : SNCA +NT | 0.3269 | -0.3164 to 0.9702 | No | ns | 0.5337 |
| WT : WT vs. SNCA : SNCA +ASO | 0.1358 | -0.5075 to 0.7791 | No | ns | 0.9424 |
| SNCA : SNCA +NT vs. SNCA : SNCA +ASO | -0.1911 | -0.8344 to 0.4522 | No | ns | 0.8574 |
| Hour 6 |  |  |  |  |  |
| WT : WT vs. SNCA : SNCA +NT | 0.4977 | -0.1456 to 1.141 | No | ns | 0.1814 |
| WT : WT vs. SNCA : SNCA +ASO | 0.2904 | -0.3529 to 0.9337 | No | ns | 0.6275 |
| SNCA : SNCA +NT vs. SNCA : SNCA +ASO | -0.2073 | -0.8506 to 0.4360 | No | ns | 0.8254 |
| Hour 7 |  |  |  |  |  |
| WT : WT vs. SNCA : SNCA +NT | 0.6603 | 0.01696 to 1.304 | Yes | * | 0.0422 |
| WT : WT vs. SNCA : SNCA +ASO | 0.4367 | -0.2066 to 1.080 | No | ns | 0.2825 |
| SNCA : SNCA +NT vs. SNCA : SNCA +ASO | -0.2235 | -0.8669 to 0.4198 | No | ns | 0.7905 |
| Hour 8 |  |  |  |  |  |
| WT : WT vs. SNCA : SNCA +NT | 0.9517 | 0.3084 to 1.595 | Yes | ** | 0.0013 |
| WT : WT vs. SNCA : SNCA +ASO | 0.7017 | 0.05834 to 1.345 | Yes | * | 0.0274 |
| SNCA : SNCA +NT vs. SNCA : SNCA +ASO | -0.25 | -0.8933 to 0.3933 | No | ns | 0.7289 |
| Hour 9 |  |  |  |  |  |
| WT : WT vs. SNCA : SNCA +NT | 1.155 | 0.5113 to 1.798 | Yes | **** | <0.0001 |
| WT : WT vs. SNCA : SNCA +ASO | 0.8656 | 0.2222 to 1.509 | Yes | ** | 0.004 |
| SNCA : SNCA +NT vs. SNCA : SNCA +ASO | -0.289 | -0.9323 to 0.3543 | No | ns | 0.6312 |
| Hour 10 |  |  |  |  |  |
| WT : WT vs. SNCA : SNCA +NT | 1.275 | 0.6319 to 1.919 | Yes | **** | <0.0001 |
| WT : WT vs. SNCA : SNCA +ASO | 0.9489 | 0.3056 to 1.592 | Yes | ** | 0.0013 |
| SNCA : SNCA +NT vs. SNCA : SNCA +ASO | -0.3263 | -0.9696 to 0.3171 | No | ns | 0.5353 |
| Hour 11 |  |  |  |  |  |
| WT : WT vs. SNCA : SNCA +NT | 1.308 | 0.6649 to 1.952 | Yes | **** | <0.0001 |
| WT : WT vs. SNCA : SNCA +ASO | 0.9885 | 0.3452 to 1.632 | Yes | *** | 0.0008 |
| SNCA : SNCA +NT vs. SNCA : SNCA +ASO | -0.3197 | -0.9630 to 0.3236 | No | ns | 0.5522 |
| Hour 12 |  |  |  |  |  |
| WT : WT vs. SNCA : SNCA +NT | 1.405 | 0.7613 to 2.048 | Yes | **** | <0.0001 |
| WT : WT vs. SNCA : SNCA +ASO | 1.021 | 0.3779 to 1.665 | Yes | *** | 0.0005 |
| SNCA : SNCA +NT vs. SNCA : SNCA +ASO | -0.3834 | -1.027 to 0.2599 | No | ns | 0.3953 |
| Hour 13 |  |  |  |  |  |
| WT : WT vs. SNCA : SNCA +NT | 1.439 | 0.7959 to 2.083 | Yes | **** | <0.0001 |
| WT : WT vs. SNCA : SNCA +ASO | 1.068 | 0.4246 to 1.711 | Yes | *** | 0.0002 |
| SNCA : SNCA +NT vs. SNCA : SNCA +ASO | -0.3713 | -1.015 to 0.2720 | No | ns | 0.4236 |
| Hour 14 |  |  |  |  |  |
| WT : WT vs. SNCA : SNCA +NT | 1.461 | 0.8176 to 2.104 | Yes | **** | <0.0001 |
| WT : WT vs. SNCA : SNCA +ASO | 1.005 | 0.3619 to 1.649 | Yes | *** | 0.0006 |
| SNCA : SNCA +NT vs. SNCA : SNCA +ASO | -0.4557 | -1.099 to 0.1876 | No | ns | 0.2478 |
| Hour 15 |  |  |  |  |  |
| WT : WT vs. SNCA : SNCA +NT | 1.593 | 0.9495 to 2.236 | Yes | **** | <0.0001 |
| WT : WT vs. SNCA : SNCA +ASO | 1.156 | 0.5128 to 1.799 | Yes | **** | <0.0001 |
| SNCA : SNCA +NT vs. SNCA : SNCA +ASO | -0.4367 | -1.080 to 0.2067 | No | ns | 0.2826 |
| Hour 16 |  |  |  |  |  |
| WT : WT vs. SNCA : SNCA +NT | 1.779 | 1.136 to 2.423 | Yes | **** | <0.0001 |
| WT : WT vs. SNCA : SNCA +ASO | 1.176 | 0.5326 to 1.819 | Yes | **** | <0.0001 |
| SNCA : SNCA +NT vs. SNCA : SNCA +ASO | -0.6033 | -1.247 to 0.04001 | No | ns | 0.0735 |
| Hour 17 |  |  |  |  |  |
| WT : WT vs. SNCA : SNCA +NT | 1.753 | 1.109 to 2.396 | Yes | **** | <0.0001 |
| WT : WT vs. SNCA : SNCA +ASO | 1.223 | 0.5792 to 1.866 | Yes | **** | <0.0001 |
| SNCA : SNCA +NT vs. SNCA : SNCA +ASO | -0.53 | -1.173 to 0.1133 | No | ns | 0.1401 |
| Hour 18 |  |  |  |  |  |
| WT : WT vs. SNCA : SNCA +NT | 1.92 | 1.277 to 2.563 | Yes | **** | <0.0001 |
| WT : WT vs. SNCA : SNCA +ASO | 1.311 | 0.6681 to 1.955 | Yes | **** | <0.0001 |
| SNCA : SNCA +NT vs. SNCA : SNCA +ASO | -0.6086 | -1.252 to 0.03473 | No | ns | 0.0699 |
| Hour 19 |  |  |  |  |  |
| WT : WT vs. SNCA : SNCA +NT | 2.124 | 1.480 to 2.767 | Yes | **** | <0.0001 |
| WT : WT vs. SNCA : SNCA +ASO | 1.397 | 0.7537 to 2.040 | Yes | **** | <0.0001 |
| SNCA : SNCA +NT vs. SNCA : SNCA +ASO | -0.7268 | -1.370 to -0.08350 | Yes | * | 0.0209 |
| Hour 20 |  |  |  |  |  |
| WT : WT vs. SNCA : SNCA +NT | 2.117 | 1.473 to 2.760 | Yes | **** | <0.0001 |
| WT : WT vs. SNCA : SNCA +ASO | 1.337 | 0.6937 to 1.980 | Yes | **** | <0.0001 |
| SNCA : SNCA +NT vs. SNCA : SNCA +ASO | -0.7795 | -1.423 to -0.1362 | Yes | * | 0.0115 |
| Hour 21 |  |  |  |  |  |
| WT : WT vs. SNCA : SNCA +NT | 2.621 | 1.978 to 3.265 | Yes | **** | <0.0001 |
| WT : WT vs. SNCA : SNCA +ASO | 1.622 | 0.9784 to 2.265 | Yes | **** | <0.0001 |
| SNCA : SNCA +NT vs. SNCA : SNCA +ASO | -0.9995 | -1.643 to -0.3562 | Yes | *** | 0.0006 |
| Hour 22 |  |  |  |  |  |
| WT : WT vs. SNCA : SNCA +NT | 2.508 | 1.865 to 3.151 | Yes | **** | <0.0001 |
| WT : WT vs. SNCA : SNCA +ASO | 1.582 | 0.9390 to 2.226 | Yes | **** | <0.0001 |
| SNCA : SNCA +NT vs. SNCA : SNCA +ASO | -0.9259 | -1.569 to -0.2826 | Yes | ** | 0.0018 |
| Hour 23 |  |  |  |  |  |
| WT : WT vs. SNCA : SNCA +NT | 2.595 | 1.952 to 3.239 | Yes | **** | <0.0001 |
| WT : WT vs. SNCA : SNCA +ASO | 1.76 | 1.117 to 2.403 | Yes | **** | <0.0001 |
| SNCA : SNCA +NT vs. SNCA : SNCA +ASO | -0.8354 | -1.479 to -0.1921 | Yes | ** | 0.0058 |
| Hour 24 |  |  |  |  |  |
| WT : WT vs. SNCA : SNCA +NT | 2.689 | 2.045 to 3.332 | Yes | **** | <0.0001 |
| WT : WT vs. SNCA : SNCA +ASO | 1.734 | 1.091 to 2.378 | Yes | **** | <0.0001 |
| SNCA : SNCA +NT vs. SNCA : SNCA +ASO | -0.9541 | -1.597 to -0.3108 | Yes | ** | 0.0012 |
| Hour 25 |  |  |  |  |  |
| WT : WT vs. SNCA : SNCA +NT | 2.637 | 1.994 to 3.280 | Yes | **** | <0.0001 |
| WT : WT vs. SNCA : SNCA +ASO | 1.608 | 0.9649 to 2.252 | Yes | **** | <0.0001 |
| SNCA : SNCA +NT vs. SNCA : SNCA +ASO | -1.029 | -1.672 to -0.3854 | Yes | *** | 0.0004 |
| Hour 26 |  |  |  |  |  |
| WT : WT vs. SNCA : SNCA +NT | 2.864 | 2.221 to 3.507 | Yes | **** | <0.0001 |
| WT : WT vs. SNCA : SNCA +ASO | 1.796 | 1.153 to 2.440 | Yes | **** | <0.0001 |
| SNCA : SNCA +NT vs. SNCA : SNCA +ASO | -1.068 | -1.711 to -0.4243 | Yes | *** | 0.0002 |
| Hour 27 |  |  |  |  |  |
| WT : WT vs. SNCA : SNCA +NT | 2.823 | 2.180 to 3.467 | Yes | **** | <0.0001 |
| WT : WT vs. SNCA : SNCA +ASO | 1.717 | 1.074 to 2.360 | Yes | **** | <0.0001 |
| SNCA : SNCA +NT vs. SNCA : SNCA +ASO | -1.106 | -1.750 to -0.4631 | Yes | *** | 0.0001 |
| Hour 28 |  |  |  |  |  |
| WT : WT vs. SNCA : SNCA +NT | 2.773 | 2.130 to 3.417 | Yes | **** | <0.0001 |
| WT : WT vs. SNCA : SNCA +ASO | 1.665 | 1.022 to 2.309 | Yes | **** | <0.0001 |
| SNCA : SNCA +NT vs. SNCA : SNCA +ASO | -1.108 | -1.751 to -0.4648 | Yes | *** | 0.0001 |
| Hour 29 |  |  |  |  |  |
| WT : WT vs. SNCA : SNCA +NT | 2.808 | 2.165 to 3.451 | Yes | **** | <0.0001 |
| WT : WT vs. SNCA : SNCA +ASO | 1.677 | 1.034 to 2.321 | Yes | **** | <0.0001 |
| SNCA : SNCA +NT vs. SNCA : SNCA +ASO | -1.131 | -1.774 to -0.4875 | Yes | **** | <0.0001 |
| Hour 30 |  |  |  |  |  |
| WT : WT vs. SNCA : SNCA +NT | 3.024 | 2.381 to 3.667 | Yes | **** | <0.0001 |
| WT : WT vs. SNCA : SNCA +ASO | 1.82 | 1.176 to 2.463 | Yes | **** | <0.0001 |
| SNCA : SNCA +NT vs. SNCA : SNCA +ASO | -1.204 | -1.847 to -0.5607 | Yes | **** | <0.0001 |
| Hour 31 |  |  |  |  |  |
| WT : WT vs. SNCA : SNCA +NT | 2.876 | 2.232 to 3.519 | Yes | **** | <0.0001 |
| WT : WT vs. SNCA : SNCA +ASO | 1.74 | 1.097 to 2.384 | Yes | **** | <0.0001 |
| SNCA : SNCA +NT vs. SNCA : SNCA +ASO | -1.135 | -1.779 to -0.4921 | Yes | **** | <0.0001 |
| Hour 32 |  |  |  |  |  |
| WT : WT vs. SNCA : SNCA +NT | 2.902 | 2.259 to 3.545 | Yes | **** | <0.0001 |
| WT : WT vs. SNCA : SNCA +ASO | 1.702 | 1.058 to 2.345 | Yes | **** | <0.0001 |
| SNCA : SNCA +NT vs. SNCA : SNCA +ASO | -1.2 | -1.844 to -0.5569 | Yes | **** | <0.0001 |
| Hour 33 |  |  |  |  |  |
| WT : WT vs. SNCA : SNCA +NT | 3.006 | 2.362 to 3.649 | Yes | **** | <0.0001 |
| WT : WT vs. SNCA : SNCA +ASO | 1.692 | 1.048 to 2.335 | Yes | **** | <0.0001 |
| SNCA : SNCA +NT vs. SNCA : SNCA +ASO | -1.314 | -1.957 to -0.6707 | Yes | **** | <0.0001 |
| Hour 34 |  |  |  |  |  |
| WT : WT vs. SNCA : SNCA +NT | 3.135 | 2.492 to 3.778 | Yes | **** | <0.0001 |
| WT : WT vs. SNCA : SNCA +ASO | 1.755 | 1.112 to 2.398 | Yes | **** | <0.0001 |
| SNCA : SNCA +NT vs. SNCA : SNCA +ASO | -1.38 | -2.023 to -0.7366 | Yes | **** | <0.0001 |
| Hour 35 |  |  |  |  |  |
| WT : WT vs. SNCA : SNCA +NT | 3.088 | 2.445 to 3.732 | Yes | **** | <0.0001 |
| WT : WT vs. SNCA : SNCA +ASO | 1.692 | 1.049 to 2.336 | Yes | **** | <0.0001 |
| SNCA : SNCA +NT vs. SNCA : SNCA +ASO | -1.396 | -2.039 to -0.7527 | Yes | **** | <0.0001 |
| Hour 36 |  |  |  |  |  |
| WT : WT vs. SNCA : SNCA +NT | 3.109 | 2.466 to 3.753 | Yes | **** | <0.0001 |
| WT : WT vs. SNCA : SNCA +ASO | 1.632 | 0.9882 to 2.275 | Yes | **** | <0.0001 |
| SNCA : SNCA +NT vs. SNCA : SNCA +ASO | -1.478 | -2.121 to -0.8345 | Yes | **** | <0.0001 |
| Hour 37 |  |  |  |  |  |
| WT : WT vs. SNCA : SNCA +NT | 3.05 | 2.407 to 3.693 | Yes | **** | <0.0001 |
| WT : WT vs. SNCA : SNCA +ASO | 1.551 | 0.9072 to 2.194 | Yes | **** | <0.0001 |
| SNCA : SNCA +NT vs. SNCA : SNCA +ASO | -1.499 | -2.143 to -0.8561 | Yes | **** | <0.0001 |
| Hour 38 |  |  |  |  |  |
| WT : WT vs. SNCA : SNCA +NT | 2.954 | 2.311 to 3.597 | Yes | **** | <0.0001 |
| WT : WT vs. SNCA : SNCA +ASO | 1.528 | 0.8850 to 2.172 | Yes | **** | <0.0001 |
| SNCA : SNCA +NT vs. SNCA : SNCA +ASO | -1.426 | -2.069 to -0.7823 | Yes | **** | <0.0001 |
| Hour 39 |  |  |  |  |  |
| WT : WT vs. SNCA : SNCA +NT | 3.129 | 2.486 to 3.773 | Yes | **** | <0.0001 |
| WT : WT vs. SNCA : SNCA +ASO | 1.633 | 0.9901 to 2.277 | Yes | **** | <0.0001 |
| SNCA : SNCA +NT vs. SNCA : SNCA +ASO | -1.496 | -2.139 to -0.8527 | Yes | **** | <0.0001 |
| Hour 40 |  |  |  |  |  |
| WT : WT vs. SNCA : SNCA +NT | 3.33 | 2.686 to 3.973 | Yes | **** | <0.0001 |
| WT : WT vs. SNCA : SNCA +ASO | 1.656 | 1.013 to 2.299 | Yes | **** | <0.0001 |
| SNCA : SNCA +NT vs. SNCA : SNCA +ASO | -1.674 | -2.317 to -1.030 | Yes | **** | <0.0001 |
| Hour 41 |  |  |  |  |  |
| WT : WT vs. SNCA : SNCA +NT | 3.254 | 2.610 to 3.897 | Yes | **** | <0.0001 |
| WT : WT vs. SNCA : SNCA +ASO | 1.424 | 0.7802 to 2.067 | Yes | **** | <0.0001 |
| SNCA : SNCA +NT vs. SNCA : SNCA +ASO | -1.83 | -2.473 to -1.187 | Yes | **** | <0.0001 |
| Hour 42 |  |  |  |  |  |
| WT : WT vs. SNCA : SNCA +NT | 3.184 | 2.540 to 3.827 | Yes | **** | <0.0001 |
| WT : WT vs. SNCA : SNCA +ASO | 1.392 | 0.7484 to 2.035 | Yes | **** | <0.0001 |
| SNCA : SNCA +NT vs. SNCA : SNCA +ASO | -1.792 | -2.435 to -1.149 | Yes | **** | <0.0001 |
| Hour 43 |  |  |  |  |  |
| WT : WT vs. SNCA : SNCA +NT | 3.134 | 2.490 to 3.777 | Yes | **** | <0.0001 |
| WT : WT vs. SNCA : SNCA +ASO | 1.463 | 0.8194 to 2.106 | Yes | **** | <0.0001 |
| SNCA : SNCA +NT vs. SNCA : SNCA +ASO | -1.671 | -2.314 to -1.028 | Yes | **** | <0.0001 |
| Hour 44 |  |  |  |  |  |
| WT : WT vs. SNCA : SNCA +NT | 3.113 | 2.470 to 3.757 | Yes | **** | <0.0001 |
| WT : WT vs. SNCA : SNCA +ASO | 1.412 | 0.7688 to 2.055 | Yes | **** | <0.0001 |
| SNCA : SNCA +NT vs. SNCA : SNCA +ASO | -1.701 | -2.344 to -1.058 | Yes | **** | <0.0001 |
| Hour 45 |  |  |  |  |  |
| WT : WT vs. SNCA : SNCA +NT | 3.281 | 2.637 to 3.924 | Yes | **** | <0.0001 |
| WT : WT vs. SNCA : SNCA +ASO | 1.457 | 0.8132 to 2.100 | Yes | **** | <0.0001 |
| SNCA : SNCA +NT vs. SNCA : SNCA +ASO | -1.824 | -2.467 to -1.181 | Yes | **** | <0.0001 |
| Hour 46 |  |  |  |  |  |
| WT : WT vs. SNCA : SNCA +NT | 3.101 | 2.458 to 3.744 | Yes | **** | <0.0001 |
| WT : WT vs. SNCA : SNCA +ASO | 1.353 | 0.7092 to 1.996 | Yes | **** | <0.0001 |
| SNCA : SNCA +NT vs. SNCA : SNCA +ASO | -1.748 | -2.392 to -1.105 | Yes | **** | <0.0001 |
| Hour 47 |  |  |  |  |  |
| WT : WT vs. SNCA : SNCA +NT | 3.08 | 2.437 to 3.724 | Yes | **** | <0.0001 |
| WT : WT vs. SNCA : SNCA +ASO | 1.381 | 0.7379 to 2.025 | Yes | **** | <0.0001 |
| SNCA : SNCA +NT vs. SNCA : SNCA +ASO | -1.699 | -2.342 to -1.056 | Yes | **** | <0.0001 |
| Hour 48 |  |  |  |  |  |
| WT : WT vs. SNCA : SNCA +NT | 3.25 | 2.607 to 3.893 | Yes | **** | <0.0001 |
| WT : WT vs. SNCA : SNCA +ASO | 1.429 | 0.7861 to 2.073 | Yes | **** | <0.0001 |
| SNCA : SNCA +NT vs. SNCA : SNCA +ASO | -1.82 | -2.464 to -1.177 | Yes | **** | <0.0001 |

**Supplemental Table 3:** Sidak’s Multiple Comparisons Post-hoc Test of Phagocytosis Following ASO Treatment in Mixed Co-Cultures.

|  |  |  |  |  |  |
| --- | --- | --- | --- | --- | --- |
| Sidak's multiple comparisons test | Mean Diff. | 95.00% CI of diff. | Below threshold? | Summary | Adjusted P Value |
| Hour 1 |  |  |  |  |  |
| WT : WT vs. SNCA : WT +NT | 0.05839 | -0.6606 to 0.7774 | No | ns | 0.9963 |
| WT : WT vs. SNCA : WT +ASO | -0.01665 | -0.7356 to 0.7023 | No | ns | >0.9999 |
| SNCA : WT +NT vs. SNCA : WT +ASO | -0.07504 | -0.7940 to 0.6439 | No | ns | 0.9923 |
| Hour 2 |  |  |  |  |  |
| WT : WT vs. SNCA : WT +NT | 0.002212 | -0.7168 to 0.7212 | No | ns | >0.9999 |
| WT : WT vs. SNCA : WT +ASO | -0.05925 | -0.7782 to 0.6597 | No | ns | 0.9962 |
| SNCA : WT +NT vs. SNCA : WT +ASO | -0.06146 | -0.7804 to 0.6575 | No | ns | 0.9958 |
| Hour 3 |  |  |  |  |  |
| WT : WT vs. SNCA : WT +NT | -0.07979 | -0.7987 to 0.6392 | No | ns | 0.9908 |
| WT : WT vs. SNCA : WT +ASO | -0.1153 | -0.8343 to 0.6036 | No | ns | 0.9734 |
| SNCA : WT +NT vs. SNCA : WT +ASO | -0.03554 | -0.7545 to 0.6834 | No | ns | 0.9992 |
| Hour 4 |  |  |  |  |  |
| WT : WT vs. SNCA : WT +NT | -0.04755 | -0.7665 to 0.6714 | No | ns | 0.998 |
| WT : WT vs. SNCA : WT +ASO | -0.04457 | -0.7635 to 0.6744 | No | ns | 0.9984 |
| SNCA : WT +NT vs. SNCA : WT +ASO | 0.002983 | -0.7160 to 0.7219 | No | ns | >0.9999 |
| Hour 5 |  |  |  |  |  |
| WT : WT vs. SNCA : WT +NT | 0.1238 | -0.5952 to 0.8427 | No | ns | 0.9674 |
| WT : WT vs. SNCA : WT +ASO | 0.1353 | -0.5836 to 0.8543 | No | ns | 0.9581 |
| SNCA : WT +NT vs. SNCA : WT +ASO | 0.01156 | -0.7074 to 0.7305 | No | ns | >0.9999 |
| Hour 6 |  |  |  |  |  |
| WT : WT vs. SNCA : WT +NT | 0.273 | -0.4460 to 0.9919 | No | ns | 0.7428 |
| WT : WT vs. SNCA : WT +ASO | 0.2475 | -0.4714 to 0.9665 | No | ns | 0.7951 |
| SNCA : WT +NT vs. SNCA : WT +ASO | -0.02543 | -0.7444 to 0.6935 | No | ns | 0.9997 |
| Hour 7 |  |  |  |  |  |
| WT : WT vs. SNCA : WT +NT | 0.4607 | -0.2583 to 1.180 | No | ns | 0.3317 |
| WT : WT vs. SNCA : WT +ASO | 0.3818 | -0.3372 to 1.101 | No | ns | 0.4964 |
| SNCA : WT +NT vs. SNCA : WT +ASO | -0.0789 | -0.7979 to 0.6401 | No | ns | 0.9911 |
| Hour 8 |  |  |  |  |  |
| WT : WT vs. SNCA : WT +NT | 0.7219 | 0.002984 to 1.441 | Yes | * | 0.0487 |
| WT : WT vs. SNCA : WT +ASO | 0.6011 | -0.1179 to 1.320 | No | ns | 0.1313 |
| SNCA : WT +NT vs. SNCA : WT +ASO | -0.1209 | -0.8399 to 0.5981 | No | ns | 0.9695 |
| Hour 9 |  |  |  |  |  |
| WT : WT vs. SNCA : WT +NT | 0.8953 | 0.1763 to 1.614 | Yes | ** | 0.0089 |
| WT : WT vs. SNCA : WT +ASO | 0.7337 | 0.01472 to 1.453 | Yes | * | 0.0438 |
| SNCA : WT +NT vs. SNCA : WT +ASO | -0.1616 | -0.8806 to 0.5573 | No | ns | 0.9315 |
| Hour 10 |  |  |  |  |  |
| WT : WT vs. SNCA : WT +NT | 0.9462 | 0.2272 to 1.665 | Yes | ** | 0.0051 |
| WT : WT vs. SNCA : WT +ASO | 0.8103 | 0.09130 to 1.529 | Yes | * | 0.0213 |
| SNCA : WT +NT vs. SNCA : WT +ASO | -0.1359 | -0.8549 to 0.5830 | No | ns | 0.9576 |
| Hour 11 |  |  |  |  |  |
| WT : WT vs. SNCA : WT +NT | 1.077 | 0.3582 to 1.796 | Yes | ** | 0.0011 |
| WT : WT vs. SNCA : WT +ASO | 0.8424 | 0.1234 to 1.561 | Yes | * | 0.0154 |
| SNCA : WT +NT vs. SNCA : WT +ASO | -0.2348 | -0.9538 to 0.4842 | No | ns | 0.8196 |
| Hour 12 |  |  |  |  |  |
| WT : WT vs. SNCA : WT +NT | 1.203 | 0.4841 to 1.922 | Yes | *** | 0.0002 |
| WT : WT vs. SNCA : WT +ASO | 0.8856 | 0.1666 to 1.605 | Yes | ** | 0.0098 |
| SNCA : WT +NT vs. SNCA : WT +ASO | -0.3175 | -1.036 to 0.4015 | No | ns | 0.6438 |
| Hour 13 |  |  |  |  |  |
| WT : WT vs. SNCA : WT +NT | 1.262 | 0.5435 to 1.981 | Yes | **** | <0.0001 |
| WT : WT vs. SNCA : WT +ASO | 0.9097 | 0.1907 to 1.629 | Yes | ** | 0.0076 |
| SNCA : WT +NT vs. SNCA : WT +ASO | -0.3528 | -1.072 to 0.3662 | No | ns | 0.5624 |
| Hour 14 |  |  |  |  |  |
| WT : WT vs. SNCA : WT +NT | 1.277 | 0.5581 to 1.996 | Yes | **** | <0.0001 |
| WT : WT vs. SNCA : WT +ASO | 0.9269 | 0.2079 to 1.646 | Yes | ** | 0.0063 |
| SNCA : WT +NT vs. SNCA : WT +ASO | -0.3502 | -1.069 to 0.3688 | No | ns | 0.5684 |
| Hour 15 |  |  |  |  |  |
| WT : WT vs. SNCA : WT +NT | 1.433 | 0.7143 to 2.152 | Yes | **** | <0.0001 |
| WT : WT vs. SNCA : WT +ASO | 0.9979 | 0.2790 to 1.717 | Yes | ** | 0.0028 |
| SNCA : WT +NT vs. SNCA : WT +ASO | -0.4353 | -1.154 to 0.2836 | No | ns | 0.3813 |
| Hour 16 |  |  |  |  |  |
| WT : WT vs. SNCA : WT +NT | 1.477 | 0.7579 to 2.196 | Yes | **** | <0.0001 |
| WT : WT vs. SNCA : WT +ASO | 1.067 | 0.3477 to 1.786 | Yes | ** | 0.0012 |
| SNCA : WT +NT vs. SNCA : WT +ASO | -0.4103 | -1.129 to 0.3087 | No | ns | 0.4336 |
| Hour 17 |  |  |  |  |  |
| WT : WT vs. SNCA : WT +NT | 1.413 | 0.6940 to 2.132 | Yes | **** | <0.0001 |
| WT : WT vs. SNCA : WT +ASO | 1.016 | 0.2970 to 1.735 | Yes | ** | 0.0023 |
| SNCA : WT +NT vs. SNCA : WT +ASO | -0.397 | -1.116 to 0.3219 | No | ns | 0.4624 |
| Hour 18 |  |  |  |  |  |
| WT : WT vs. SNCA : WT +NT | 1.551 | 0.8324 to 2.270 | Yes | **** | <0.0001 |
| WT : WT vs. SNCA : WT +ASO | 1.085 | 0.3656 to 1.804 | Yes | *** | 0.001 |
| SNCA : WT +NT vs. SNCA : WT +ASO | -0.4668 | -1.186 to 0.2521 | No | ns | 0.3202 |
| Hour 19 |  |  |  |  |  |
| WT : WT vs. SNCA : WT +NT | 1.637 | 0.9182 to 2.356 | Yes | **** | <0.0001 |
| WT : WT vs. SNCA : WT +ASO | 1.115 | 0.3961 to 1.834 | Yes | *** | 0.0007 |
| SNCA : WT +NT vs. SNCA : WT +ASO | -0.5221 | -1.241 to 0.1969 | No | ns | 0.2282 |
| Hour 20 |  |  |  |  |  |
| WT : WT vs. SNCA : WT +NT | 1.604 | 0.8847 to 2.323 | Yes | **** | <0.0001 |
| WT : WT vs. SNCA : WT +ASO | 0.9848 | 0.2658 to 1.704 | Yes | ** | 0.0033 |
| SNCA : WT +NT vs. SNCA : WT +ASO | -0.6189 | -1.338 to 0.1001 | No | ns | 0.1146 |
| Hour 21 |  |  |  |  |  |
| WT : WT vs. SNCA : WT +NT | 1.888 | 1.169 to 2.607 | Yes | **** | <0.0001 |
| WT : WT vs. SNCA : WT +ASO | 1.115 | 0.3964 to 1.834 | Yes | *** | 0.0007 |
| SNCA : WT +NT vs. SNCA : WT +ASO | -0.7722 | -1.491 to -0.05325 | Yes | * | 0.0307 |
| Hour 22 |  |  |  |  |  |
| WT : WT vs. SNCA : WT +NT | 1.875 | 1.156 to 2.594 | Yes | **** | <0.0001 |
| WT : WT vs. SNCA : WT +ASO | 1.041 | 0.3225 to 1.760 | Yes | ** | 0.0017 |
| SNCA : WT +NT vs. SNCA : WT +ASO | -0.8338 | -1.553 to -0.1148 | Yes | * | 0.0168 |
| Hour 23 |  |  |  |  |  |
| WT : WT vs. SNCA : WT +NT | 1.955 | 1.236 to 2.674 | Yes | **** | <0.0001 |
| WT : WT vs. SNCA : WT +ASO | 1.104 | 0.3847 to 1.823 | Yes | *** | 0.0008 |
| SNCA : WT +NT vs. SNCA : WT +ASO | -0.8514 | -1.570 to -0.1325 | Yes | * | 0.0141 |
| Hour 24 |  |  |  |  |  |
| WT : WT vs. SNCA : WT +NT | 1.816 | 1.097 to 2.535 | Yes | **** | <0.0001 |
| WT : WT vs. SNCA : WT +ASO | 0.9796 | 0.2606 to 1.699 | Yes | ** | 0.0035 |
| SNCA : WT +NT vs. SNCA : WT +ASO | -0.8363 | -1.555 to -0.1173 | Yes | * | 0.0164 |
| Hour 25 |  |  |  |  |  |
| WT : WT vs. SNCA : WT +NT | 1.768 | 1.049 to 2.487 | Yes | **** | <0.0001 |
| WT : WT vs. SNCA : WT +ASO | 0.8647 | 0.1457 to 1.584 | Yes | * | 0.0123 |
| SNCA : WT +NT vs. SNCA : WT +ASO | -0.9034 | -1.622 to -0.1844 | Yes | ** | 0.0081 |
| Hour 26 |  |  |  |  |  |
| WT : WT vs. SNCA : WT +NT | 1.922 | 1.203 to 2.641 | Yes | **** | <0.0001 |
| WT : WT vs. SNCA : WT +ASO | 0.8901 | 0.1711 to 1.609 | Yes | ** | 0.0094 |
| SNCA : WT +NT vs. SNCA : WT +ASO | -1.032 | -1.751 to -0.3130 | Yes | ** | 0.0019 |
| Hour 27 |  |  |  |  |  |
| WT : WT vs. SNCA : WT +NT | 1.784 | 1.065 to 2.503 | Yes | **** | <0.0001 |
| WT : WT vs. SNCA : WT +ASO | 0.863 | 0.1440 to 1.582 | Yes | * | 0.0125 |
| SNCA : WT +NT vs. SNCA : WT +ASO | -0.9207 | -1.640 to -0.2018 | Yes | ** | 0.0067 |
| Hour 28 |  |  |  |  |  |
| WT : WT vs. SNCA : WT +NT | 1.774 | 1.055 to 2.493 | Yes | **** | <0.0001 |
| WT : WT vs. SNCA : WT +ASO | 0.7407 | 0.02175 to 1.460 | Yes | * | 0.0411 |
| SNCA : WT +NT vs. SNCA : WT +ASO | -1.033 | -1.752 to -0.3142 | Yes | ** | 0.0018 |
| Hour 29 |  |  |  |  |  |
| WT : WT vs. SNCA : WT +NT | 1.945 | 1.226 to 2.664 | Yes | **** | <0.0001 |
| WT : WT vs. SNCA : WT +ASO | 0.772 | 0.05299 to 1.491 | Yes | * | 0.0308 |
| SNCA : WT +NT vs. SNCA : WT +ASO | -1.173 | -1.892 to -0.4544 | Yes | *** | 0.0003 |
| Hour 30 |  |  |  |  |  |
| WT : WT vs. SNCA : WT +NT | 2.16 | 1.441 to 2.879 | Yes | **** | <0.0001 |
| WT : WT vs. SNCA : WT +ASO | 0.8391 | 0.1202 to 1.558 | Yes | * | 0.016 |
| SNCA : WT +NT vs. SNCA : WT +ASO | -1.321 | -2.040 to -0.6021 | Yes | **** | <0.0001 |
| Hour 31 |  |  |  |  |  |
| WT : WT vs. SNCA : WT +NT | 1.954 | 1.235 to 2.673 | Yes | **** | <0.0001 |
| WT : WT vs. SNCA : WT +ASO | 0.822 | 0.1030 to 1.541 | Yes | * | 0.019 |
| SNCA : WT +NT vs. SNCA : WT +ASO | -1.132 | -1.851 to -0.4127 | Yes | *** | 0.0005 |
| Hour 32 |  |  |  |  |  |
| WT : WT vs. SNCA : WT +NT | 1.93 | 1.211 to 2.649 | Yes | **** | <0.0001 |
| WT : WT vs. SNCA : WT +ASO | 0.6332 | -0.08572 to 1.352 | No | ns | 0.1025 |
| SNCA : WT +NT vs. SNCA : WT +ASO | -1.297 | -2.016 to -0.5779 | Yes | **** | <0.0001 |
| Hour 33 |  |  |  |  |  |
| WT : WT vs. SNCA : WT +NT | 2.034 | 1.316 to 2.753 | Yes | **** | <0.0001 |
| WT : WT vs. SNCA : WT +ASO | 0.691 | -0.02794 to 1.410 | No | ns | 0.0638 |
| SNCA : WT +NT vs. SNCA : WT +ASO | -1.343 | -2.062 to -0.6245 | Yes | **** | <0.0001 |
| Hour 34 |  |  |  |  |  |
| WT : WT vs. SNCA : WT +NT | 2.205 | 1.486 to 2.924 | Yes | **** | <0.0001 |
| WT : WT vs. SNCA : WT +ASO | 0.6809 | -0.03804 to 1.400 | No | ns | 0.0695 |
| SNCA : WT +NT vs. SNCA : WT +ASO | -1.524 | -2.243 to -0.8054 | Yes | **** | <0.0001 |
| Hour 35 |  |  |  |  |  |
| WT : WT vs. SNCA : WT +NT | 2.15 | 1.431 to 2.869 | Yes | **** | <0.0001 |
| WT : WT vs. SNCA : WT +ASO | 0.5456 | -0.1734 to 1.265 | No | ns | 0.1952 |
| SNCA : WT +NT vs. SNCA : WT +ASO | -1.605 | -2.324 to -0.8858 | Yes | **** | <0.0001 |
| Hour 36 |  |  |  |  |  |
| WT : WT vs. SNCA : WT +NT | 2.204 | 1.485 to 2.923 | Yes | **** | <0.0001 |
| WT : WT vs. SNCA : WT +ASO | 0.6843 | -0.03462 to 1.403 | No | ns | 0.0675 |
| SNCA : WT +NT vs. SNCA : WT +ASO | -1.519 | -2.238 to -0.8005 | Yes | **** | <0.0001 |
| Hour 37 |  |  |  |  |  |
| WT : WT vs. SNCA : WT +NT | 2.185 | 1.466 to 2.904 | Yes | **** | <0.0001 |
| WT : WT vs. SNCA : WT +ASO | 0.4212 | -0.2978 to 1.140 | No | ns | 0.4104 |
| SNCA : WT +NT vs. SNCA : WT +ASO | -1.764 | -2.483 to -1.045 | Yes | **** | <0.0001 |
| Hour 38 |  |  |  |  |  |
| WT : WT vs. SNCA : WT +NT | 2.046 | 1.327 to 2.765 | Yes | **** | <0.0001 |
| WT : WT vs. SNCA : WT +ASO | 0.3427 | -0.3763 to 1.062 | No | ns | 0.5857 |
| SNCA : WT +NT vs. SNCA : WT +ASO | -1.703 | -2.422 to -0.9843 | Yes | **** | <0.0001 |
| Hour 39 |  |  |  |  |  |
| WT : WT vs. SNCA : WT +NT | 2.229 | 1.510 to 2.948 | Yes | **** | <0.0001 |
| WT : WT vs. SNCA : WT +ASO | 0.4902 | -0.2288 to 1.209 | No | ns | 0.2789 |
| SNCA : WT +NT vs. SNCA : WT +ASO | -1.739 | -2.458 to -1.020 | Yes | **** | <0.0001 |
| Hour 40 |  |  |  |  |  |
| WT : WT vs. SNCA : WT +NT | 2.283 | 1.564 to 3.002 | Yes | **** | <0.0001 |
| WT : WT vs. SNCA : WT +ASO | 0.4025 | -0.3165 to 1.121 | No | ns | 0.4504 |
| SNCA : WT +NT vs. SNCA : WT +ASO | -1.881 | -2.600 to -1.162 | Yes | **** | <0.0001 |
| Hour 41 |  |  |  |  |  |
| WT : WT vs. SNCA : WT +NT | 2.293 | 1.574 to 3.012 | Yes | **** | <0.0001 |
| WT : WT vs. SNCA : WT +ASO | 0.3755 | -0.3435 to 1.094 | No | ns | 0.5105 |
| SNCA : WT +NT vs. SNCA : WT +ASO | -1.917 | -2.636 to -1.198 | Yes | **** | <0.0001 |
| Hour 42 |  |  |  |  |  |
| WT : WT vs. SNCA : WT +NT | 2.238 | 1.519 to 2.957 | Yes | **** | <0.0001 |
| WT : WT vs. SNCA : WT +ASO | 0.2632 | -0.4558 to 0.9822 | No | ns | 0.7634 |
| SNCA : WT +NT vs. SNCA : WT +ASO | -1.975 | -2.694 to -1.256 | Yes | **** | <0.0001 |
| Hour 43 |  |  |  |  |  |
| WT : WT vs. SNCA : WT +NT | 2.276 | 1.557 to 2.995 | Yes | **** | <0.0001 |
| WT : WT vs. SNCA : WT +ASO | 0.3595 | -0.3594 to 1.078 | No | ns | 0.547 |
| SNCA : WT +NT vs. SNCA : WT +ASO | -1.916 | -2.635 to -1.197 | Yes | **** | <0.0001 |
| Hour 44 |  |  |  |  |  |
| WT : WT vs. SNCA : WT +NT | 2.314 | 1.595 to 3.033 | Yes | **** | <0.0001 |
| WT : WT vs. SNCA : WT +ASO | 0.2718 | -0.4472 to 0.9907 | No | ns | 0.7454 |
| SNCA : WT +NT vs. SNCA : WT +ASO | -2.042 | -2.761 to -1.324 | Yes | **** | <0.0001 |
| Hour 45 |  |  |  |  |  |
| WT : WT vs. SNCA : WT +NT | 2.392 | 1.673 to 3.111 | Yes | **** | <0.0001 |
| WT : WT vs. SNCA : WT +ASO | 0.3959 | -0.3230 to 1.115 | No | ns | 0.4648 |
| SNCA : WT +NT vs. SNCA : WT +ASO | -1.996 | -2.715 to -1.277 | Yes | **** | <0.0001 |
| Hour 46 |  |  |  |  |  |
| WT : WT vs. SNCA : WT +NT | 2.332 | 1.613 to 3.051 | Yes | **** | <0.0001 |
| WT : WT vs. SNCA : WT +ASO | 0.3432 | -0.3758 to 1.062 | No | ns | 0.5846 |
| SNCA : WT +NT vs. SNCA : WT +ASO | -1.989 | -2.708 to -1.270 | Yes | **** | <0.0001 |
| Hour 47 |  |  |  |  |  |
| WT : WT vs. SNCA : WT +NT | 2.324 | 1.605 to 3.043 | Yes | **** | <0.0001 |
| WT : WT vs. SNCA : WT +ASO | 0.2984 | -0.4206 to 1.017 | No | ns | 0.6871 |
| SNCA : WT +NT vs. SNCA : WT +ASO | -2.026 | -2.745 to -1.307 | Yes | **** | <0.0001 |
| Hour 48 |  |  |  |  |  |
| WT : WT vs. SNCA : WT +NT | 2.606 | 1.887 to 3.325 | Yes | **** | <0.0001 |
| WT : WT vs. SNCA : WT +ASO | 0.5365 | -0.1825 to 1.255 | No | ns | 0.2076 |
| SNCA : WT +NT vs. SNCA : WT +ASO | -2.07 | -2.789 to -1.351 | Yes | **** | <0.0001 |

## References

1. Spillantini MG, Crowther RA, Jakes R, Hasegawa M, Goedert M. alpha-Synuclein in filamentous inclusions of Lewy bodies from Parkinson’s disease and dementia with lewy bodies. Proc Natl Acad Sci U S A. 1998;95(11):6469–73.

2. Okazaki H, Lipkin LE, Aronson SM. Diffuse intracytoplasmic ganglionic inclusions (Lewy type) associated with progressive dementia and quadriparesis in flexion. J Neuropathol Exp Neurol. 1961;20:237–44.

3. Klein C, Westenberger A. Genetics of Parkinson’s disease. Cold Spring Harb Perspect Med. 2012;2(1):a008888.

4. Lashuel HA, Overk CR, Oueslati A, Masliah E. The many faces of alpha-synuclein: from structure and toxicity to therapeutic target. Nat Rev Neurosci. 2013;14(1):38–48.

5. Blauwendraat C, Heilbron K, Vallerga CL, Bandres-Ciga S, von Coelln R, Pihlstrom L, et al. Parkinson’s disease age at onset genome-wide association study: Defining heritability, genetic loci, and alpha-synuclein mechanisms. Mov Disord. 2019;34(6):866–75.

6. Singleton AB, Farrer M, Johnson J, Singleton A, Hague S, Kachergus J, et al. alpha-Synuclein locus triplication causes Parkinson’s disease. Science. 2003;302(5646):841.

7. Ibanez P, Bonnet AM, Debarges B, Lohmann E, Tison F, Pollak P, et al. Causal relation between alpha-synuclein gene duplication and familial Parkinson’s disease. Lancet. 2004;364(9440):1169–71.

8. Chartier-Harlin MC, Kachergus J, Roumier C, Mouroux V, Douay X, Lincoln S, et al. Alpha-synuclein locus duplication as a cause of familial Parkinson’s disease. Lancet. 2004;364(9440):1167–9.

9. Nuytemans K, Theuns J, Cruts M, Van Broeckhoven C. Genetic etiology of Parkinson disease associated with mutations in the SNCA, PARK2, PINK1, PARK7, and LRRK2 genes: a mutation update. Hum Mutat. 2010;31(7):763–80.

10. Devine MJ, Ryten M, Vodicka P, Thomson AJ, Burdon T, Houlden H, et al. Parkinson’s disease induced pluripotent stem cells with triplication of the alpha-synuclein locus. Nat Commun. 2011;2:440.

11. Rosborough K, Patel N, Kalia LV. alpha-Synuclein and Parkinsonism: Updates and Future Perspectives. Curr Neurol Neurosci Rep. 2017;17(4):31.

12. Yin J, Valin KL, Dixon ML, Leavenworth JW. The Role of Microglia and Macrophages in CNS Homeostasis, Autoimmunity, and Cancer. J Immunol Res. 2017;2017:5150678.

13. Janda E, Boi L, Carta AR. Microglial Phagocytosis and Its Regulation: A Therapeutic Target in Parkinson’s Disease? Front Mol Neurosci. 2018;11:144.

14. Gardai SJ, Mao W, Schule B, Babcock M, Schoebel S, Lorenzana C, et al. Elevated alpha-synuclein impairs innate immune cell function and provides a potential peripheral biomarker for Parkinson’s disease. PLoS One. 2013;8(8):e71634.

15. Haenseler W, Zambon F, Lee H, Vowles J, Rinaldi F, Duggal G, et al. Excess alpha-synuclein compromises phagocytosis in iPSC-derived macrophages. Sci Rep. 2017;7(1):9003.

16. Rojanathammanee L, Murphy EJ, Combs CK. Expression of mutant alpha-synuclein modulates microglial phenotype in vitro. J Neuroinflammation. 2011;8:44.

17. Schafer DP, Lehrman EK, Kautzman AG, Koyama R, Mardinly AR, Yamasaki R, et al. Microglia sculpt postnatal neural circuits in an activity and complement-dependent manner. Neuron. 2012;74(4):691–705.

18. Zhan Y, Paolicelli RC, Sforazzini F, Weinhard L, Bolasco G, Pagani F, et al. Deficient neuron-microglia signaling results in impaired functional brain connectivity and social behavior. Nat Neurosci. 2014;17(3):400–6.

19. Paolicelli RC, Bolasco G, Pagani F, Maggi L, Scianni M, Panzanelli P, et al. Synaptic pruning by microglia is necessary for normal brain development. Science. 2011;333(6048):1456–8.

20. Haenseler W, Sansom SN, Buchrieser J, Newey SE, Moore CS, Nicholls FJ, et al. A Highly Efficient Human Pluripotent Stem Cell Microglia Model Displays a Neuronal-Co-culture-Specific Expression Profile and Inflammatory Response. Stem Cell Reports. 2017;8(6):1727–42.

21. Reich M, Paris I, Ebeling M, Dahm N, Schweitzer C, Reinhardt D, et al. Alzheimer’s Risk Gene TREM2 Determines Functional Properties of New Type of Human iPSC-Derived Microglia. Front Immunol. 2020;11:617860.

22. Takahashi K, Tanabe K, Ohnuki M, Narita M, Ichisaka T, Tomoda K, et al. Induction of pluripotent stem cells from adult human fibroblasts by defined factors. Cell. 2007;131(5):861–72.

23. Mertens J, Marchetto MC, Bardy C, Gage FH. Evaluating cell reprogramming, differentiation and conversion technologies in neuroscience. Nat Rev Neurosci. 2016;17(7):424–37.

24. Zhang Y, Pak C, Han Y, Ahlenius H, Zhang Z, Chanda S, et al. Rapid single-step induction of functional neurons from human pluripotent stem cells. Neuron. 2013;78(5):785–98.

25. Abud EM, Ramirez RN, Martinez ES, Healy LM, Nguyen CHH, Newman Sa, et al. iPSC-Derived Human Microglia-like Cells to Study Neurological Diseases. Neuron. 2017;94(2):278–93 e9.

26. Douvaras P, Sun B, Wang M, Kruglikov I, Lallos G, Zimmer M, et al. Directed Differentiation of Human Pluripotent Stem Cells to Microglia. Stem Cell Reports. 2017;8(6):1516–24.

27. Muffat J, Li Y, Yuan B, Mitalipova M, Omer A, Corcoran S, et al. Efficient derivation of microglia-like cells from human pluripotent stem cells. Nat Med. 2016;22(11):1358–67.

28. van Wilgenburg B, Browne C, Vowles J, Cowley SA. Efficient, long term production of monocyte-derived macrophages from human pluripotent stem cells under partly-defined and fully-defined conditions. PLoS One. 2013;8(8):e71098.

29. Graef JD, Wu H, Ng C, Sun C, Villegas V, Qadir D, et al. Partial FMRP expression is sufficient to normalize neuronal hyperactivity in Fragile X neurons. Eur J Neurosci. 2020;51(10):2143–57.

30. Tcw J, Wang M, Pimenova AA, Bowles KR, Hartley BJ, Lacin E, et al. An Efficient Platform for Astrocyte Differentiation from Human Induced Pluripotent Stem Cells. Stem Cell Reports. 2017;9(2):600–14.

31. Kapellos TS, Taylor L, Lee H, Cowley SA, James WS, Iqbal AJ, et al. A novel real time imaging platform to quantify macrophage phagocytosis. Biochem Pharmacol. 2016;116:107–19.

32. Lecours C, Bordeleau M, Cantin L, Parent M, Paolo TD, Tremblay ME. Microglial Implication in Parkinson’s Disease: Loss of Beneficial Physiological Roles or Gain of Inflammatory Functions? Front Cell Neurosci. 2018;12:282.

33. Ferreira SA, Romero-Ramos M. Microglia Response During Parkinson’s Disease: Alpha-Synuclein Intervention. Front Cell Neurosci. 2018;12:247.

34. Tagliafierro L, Chiba-Falek O. Up-regulation of SNCA gene expression: implications to synucleinopathies. Neurogenetics. 2016;17(3):145–57.

35. Booms A, Coetzee GA. Functions of Intracellular Alpha-Synuclein in Microglia: Implications for Parkinson’s Disease Risk. Front Cell Neurosci. 2021;15:759571.

36. Zharikov A, Bai Q, De Miranda BR, Van Laar A, Greenamyre JT, Burton EA. Long-term RNAi knockdown of alpha-synuclein in the adult rat substantia nigra without neurodegeneration. Neurobiol Dis. 2019;125:146–53.

37. Mittal S, Bjornevik K, Im DS, Flierl A, Dong X, Locascio JJ, et al. beta2-Adrenoreceptor is a regulator of the alpha-synuclein gene driving risk of Parkinson’s disease. Science. 2017;357(6354):891–8.

38. Menon S, Kofoed RH, Nabbouh F, Xhima K, Al-Fahoum Y, Langman T, et al. Viral alpha-synuclein knockdown prevents spreading synucleinopathy. Brain Commun. 2021;3(4):fcab247.

39. Lewis J, Melrose H, Bumcrot D, Hope A, Zehr C, Lincoln S, et al. In vivo silencing of alpha-synuclein using naked siRNA. Mol Neurodegener. 2008;3:19.

40. Zharikov AD, Cannon JR, Tapias V, Bai Q, Horowitz MP, Shah V, et al. shRNA targeting alpha-synuclein prevents neurodegeneration in a Parkinson’s disease model. J Clin Invest. 2015;125(7):2721–35.

41. Alarcon-Aris D, Recasens A, Galofre M, Carballo-Carbajal I, Zacchi N, Ruiz-Bronchal E, et al. Selective alpha-Synuclein Knockdown in Monoamine Neurons by Intranasal Oligonucleotide Delivery: Potential Therapy for Parkinson’s Disease. Mol Ther. 2018;26(2):550–67.

42. Allen Reish HE, Standaert DG. Role of alpha-synuclein in inducing innate and adaptive immunity in Parkinson disease. J Parkinsons Dis. 2015;5(1):1–19.

43. Chen Z, Trapp BD. Microglia and neuroprotection. J Neurochem. 2016;136 Suppl 1:10–7.

44. Tremblay ME, Cookson MR, Civiero L. Glial phagocytic clearance in Parkinson’s disease. Mol Neurodegener. 2019;14(1):16.

45. Ulland TK, Colonna M. TREM2 - a key player in microglial biology and Alzheimer disease. Nat Rev Neurol. 2018;14(11):667–75.

46. Garcia-Reitboeck P, Phillips A, Piers TM, Villegas-Llerena C, Butler M, Mallach A, et al. Human Induced Pluripotent Stem Cell-Derived Microglia-Like Cells Harboring TREM2 Missense Mutations Show Specific Deficits in Phagocytosis. Cell Rep. 2018;24(9):2300–11.

47. Rayaprolu S, Mullen B, Baker M, Lynch T, Finger E, Seeley WW, et al. TREM2 in neurodegeneration: evidence for association of the p.R47H variant with frontotemporal dementia and Parkinson’s disease. Mol Neurodegener. 2013;8:19.

48. Guo Y, Wei X, Yan H, Qin Y, Yan S, Liu J, et al. TREM2 deficiency aggravates alpha-synuclein-induced neurodegeneration and neuroinflammation in Parkinson’s disease models. FASEB J. 2019;33(11):12164–74.

49. Lecca D, Janda E, Mulas G, Diana A, Martino C, Angius F, et al. Boosting phagocytosis and anti-inflammatory phenotype in microglia mediates neuroprotection by PPARgamma agonist MDG548 in Parkinson’s disease models. Br J Pharmacol. 2018;175(16):3298–314.

50. Lin HC, He Z, Ebert S, Schornig M, Santel M, Nikolova MT, et al. NGN2 induces diverse neuron types from human pluripotency. Stem Cell Reports. 2021;16(9):2118–27.

